# Sharp wave-ripple clusters enhance hippocampal-neocortical engagement for memory consolidation

**DOI:** 10.64898/2026.03.27.714843

**Authors:** Mihály Vöröslakos, Christopher Lafferty, ZheYang Zheng, Nicholas Paleologos, Elisa Chinigo, Kathryn McClain, Deren Aykan, Euisik Yoon, György Buzsáki

**Author notes:** These authors contributed equally to this work (Mihály Vöröslakos, Christopher Lafferty, ZheYang Zheng).

## Abstract

Hippocampal sharp wave-ripples (SPW-Rs), neocortical slow oscillations, and thalamocortical sleep spindles are hypothesized to provide a temporal framework for coordinated information transfer during memory consolidation. Hippocampal replay supports this process, yet replayed sequences often unfold across multiple SPW-Rs, suggesting that individual ripples may not constitute the fundamental unit of hippocampal output. Here, using large-scale electrophysiological recordings from the hippocampus and retrosplenial cortex, we show that hippocampal output is organized into clusters of SPW-Rs (cSPW-Rs) during UP states, which are often phase-locked to spindle troughs. Extending this approach with wide-field imaging and unsupervised latent-variable modeling, we found that cSPW-Rs enhanced segregation between the default mode and somatomotor networks and preferentially replayed spatially extended maze trajectories following learning. We propose that SPW-R clusters enable reverberating hippocampal-cortical spike exchange and the concatenation of sequential experiences, establishing ripple clusters as a previously unrecognized syntactic unit of hippocampal-neocortical dialogue.

## Introduction

During sleep, the brain autonomously restructures network connectivity to support the consolidation of new memories. This process is supported by a hippocampal-neocortical dialogue, thought to rely on coordinated interactions among hippocampal sharp wave-ripples (SPW-Rs), neocortical slow oscillations, and thalamocortical sleep spindles^1,2,11,3–10^. These rhythms provide a temporal framework that aligns hippocampal output with cortical depolarization, creating windows for coordinated spike transfer and synaptic plasticity^2,3,17–24,5,9,11–16^. A striking observation, however, is that the duration of hippocampal replay is often longer than that of SPW-Rs^4,12,32,23,25–31^. Consequently, many replayed neuronal sequences unfold as continuous, uninterrupted events across ripple boundaries, spanning multiple detected SPW-Rs^25,28,30,33–36^.

This mismatch in timescales suggests that the single SPW-R may not represent the unit of hippocampal output for consolidation. Instead, temporally clustered SPW-Rs may define a fundamental syntax of hippocampal-neocortical dialogue. Since ripples are commonly treated as discrete events, the defining properties and functional significance of such clustered SPW-Rs have never been directly described. Moreover, if ripple clusters constitute an effective unit of information transfer, their influence should be expressed as coordinated changes across distributed cortical systems. Whether these events preferentially reorganize functional cortical networks remains unknown. To address these questions, we combined high-density electrophysiology with widefield calcium imaging and unsupervised latent-variable modeling to show that clustered ripples broadcast replay of novel experiences while reshaping large-scale cortical dynamics for consolidation.

## Results

### Clustered SPW-Rs are prominent during NREM sleep

To characterize temporally clustered SPW-Rs, we recorded simultaneously from hippocampal CA1-CA3 regions and retrosplenial cortex (RSC), a major hippocampal output region, using four-shank Neuropixels probes^37^ and 128-channel silicon probes (**Figure 1A, B**). Inter-SPW-R intervals had a non-uniform distribution with a prominent peak at 122.5 ms (full width at half maximum = 78-182 ms) followed by a long tail (**Figure 1C**, n_SPW-Rs_ =86354 across 25 sessions in 7 mice). Double Gaussian decomposition in logarithmic space revealed two components of the interval distribution: a fast component capturing temporally clustered SPW-Rs, and a slow component representing isolated events (**Figure 1C**, bottom, **Figure S1A**). These components intersected at 177 ms, providing a boundary between clusters and isolated ripples. Notably, the short-interval peak of this fast component (126 ms) was preserved across dorsal, intermediate, and ventral recordings despite regional differences in excitability (**Figure S1B**, n_dorsal SPW-Rs_= 44439 and n_intermediate/ventral SPW-Rs_=94343, in 3 rats), indicating that ripple clusters are a conserved physiological feature along the hippocampal long axis.

**Figure 1.**
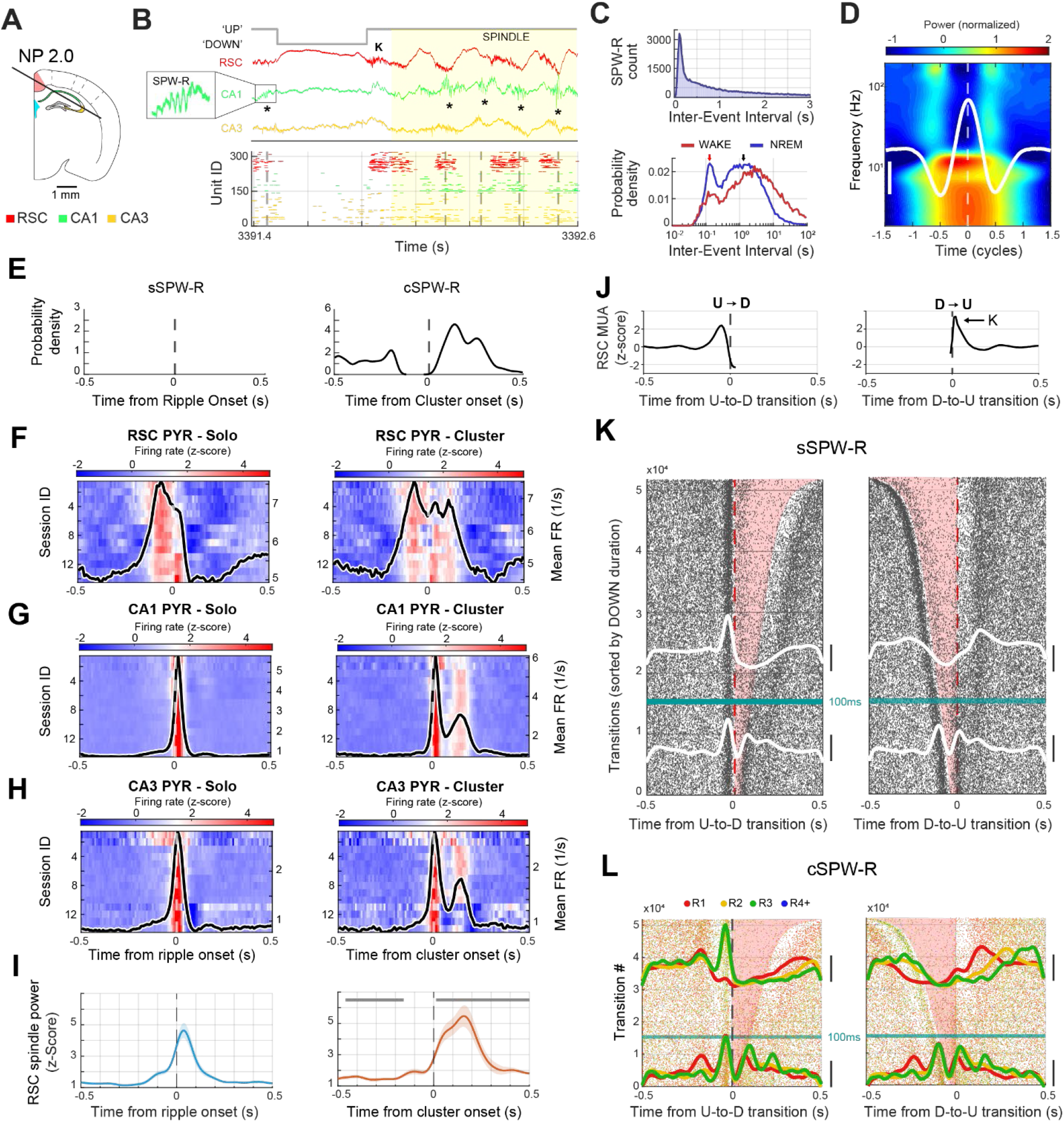
Hippocampal ripple clusters are embedded in sustained thalamocortical oscillations while solo ripples show transient cortical coupling. **(A)** Coronal schematic showing the Neuropixels 2.0 probe trajectory. Highlighted regions indicate granular RSC (red), CA1 (green), and CA3 (yellow). **(B)** Wideband LFP traces from RSC (red), CA1 (green), and CA3 (yellow). Cortical UP-DOWN transitions (solid grey line) and spindle oscillations (yellow highlight, 5-15 Hz) were identified from RSC. CA1 SPW-Rs (asterisks/grey dotted lines) were identified in the CA1 pyramidal layer. Inset, magnified view of the first **s**SPW-R. Note that **c**SPW-R sequences preferentially occur during cortical spindle oscillations. Bottom: raster plot shows multi-unit spiking across regions. Each row represents spikes from a putative single neuron. **(C)** Histogram of inter-event intervals (IEIs) for all recorded ripples (86354 events; 25 sessions from 7 mice). Top: Linear scale shows the characteristic heavy-tailed distribution with peak density at short intervals. Bottom: Logarithmic time scale reveals bimodal structure with distinct cluster and solo ripple populations. The red arrow indicates the local peak of the short-latency “cluster” population (intervals < 1 s) at 0.137 s. The black arrow indicates the global peak representing typical solo ripple intervals at 1.387 s. Blue and red curves show kernel density estimates for wake and NREM sleep states, demonstrating state-dependent differences in ripple clustering dynamics. **(D)** Cortical LFP (RSC) centered on detected peaks within the bandpass-filtered spindle oscillation (5-15 Hz). Time 0 represents the peak of each individual oscillatory cycle (n=1000 peaks from one session). Mean LFP waveform is overlaid in white (scale bar: 50 µV). Instantaneous frequency: 9.38 ± 2.28 Hz. Time axis normalized to ±2 cycles relative to each peak to account for frequency variability across individual oscillations. **(E)** Probability density of inter-event intervals showing the likelihood of finding neighboring ripples at various time lags. Left: solo ripples are isolated by >500 ms from any neighboring event (by classification criteria). Right: Cluster ripple onsets show dense temporal clustering, with high probability of additional ripples at short latencies (∼100-200 ms), reflecting the typical inter-ripple intervals within cluster sequences. Time 0 represents the reference ripple onset. **(F-H)** Population-average firing rates (z-scored) aligned to solo ripple onset (left) and cluster onset (right) for RSC **(F)**, CA1 **(G)**, and CA3 **(H)**. Heatmaps show session-averaged responses (n=14 sessions), and black traces indicate the mean firing rate across all sessions. Within each region, putative pyramidal cells (PYR) are shown. **(I)** Spindle band power (5-15 Hz) time course for solo (blue) versus cluster (orange) ripples (spindle power was Z-scored relative to a baseline -1.0 to -0.8s relative to ripple onset). Left: solo ripples exhibit a brief, transient increase in spindle power centered at ripple onset (time 0). Right: Cluster ripples occur during sustained spindle oscillations, with elevated power beginning before the first ripple and persisting throughout the cluster sequence. Grey bars indicate time windows of significant differences between cluster and solo conditions (cluster-based permutation testing, n=5000 permutations, n=14 sessions). **(J)** RSC multi-unit activity (MUA) surrounding UP-DOWN (left) and DOWN-UP (right) transitions (n=14 sessions in 4 mice). K refers to transient rebound population synchrony at the D-U transition, K-complex or “K”. **(K-L)** Raster plots show individual ripple events aligned to cortical state transitions (time 0) with DOWN states shaded in pink. Rows represent individual transitions sorted by DOWN state duration (40-500 ms), with horizontal green line marking 100 ms duration split. Marginal density plots (overlaid lines) show temporal distribution of ripple occurrence for short (<100 ms, below green line) versus long (>100 ms, above green line) DOWN states. **(K)** Solo ripples aligned to UP-to-DOWN (n=52701 transitions, left) and DOWN-to-UP transitions (n=52865 transitions, right). **(L)** Cluster ripples aligned to UP-to-DOWN (left), and DOWN-to-UP transitions (right). Ripples are color-coded by within-cluster position: R1 (first ripple, red), R2 (gold), R3 (green), R4+ (blue). Marginal density plots (overlaid lines) are shown for R1, R2, and R3. Data from 14 NREM sessions in 4 mice.

To reliably identify these two event types, we defined clustered SPW-Rs (**c**SPW-Rs) as events separated by <180 ms, and isolated or ‘solo’ SPW-Rs (**s**SPW-Rs) as those ≥500 ms apart since these thresholds minimized cross-contamination between components (**Figure S1A**, these thresholds were selected to maximize the separation between the fast and slow inter-event interval components while minimizing cross-contamination, see Methods). By these criteria, 44.9% of ripples occurred in **s**SPW-Rs, and 31.2% were **c**SPW-Rs (23.8% doublets, 5.9% triplets, 1.6% four or more, **Figures S1F, 2**). **c**SPW-R onsets exhibited an asymmetric cross-correlogram across all SPW-Rs (**Figure 1E**), indicating a refractory period preceding **c**SPW-R occurrence. Within **c**SPW-Rs, power increased from R1 to R2 of a cluster (“potentiation”), and all cluster ripples were significantly larger than **s**SPW-Rs (**Figure S1C-E**). We also found that clustering was strongly state-dependent since the ratio of **c**SPW-Rs/**s**SPW-Rs was more than twofold higher during NREM sleep than waking (**Figure 1C**; **Figure S1G-I**).

SPW-Rs occurred mainly during the RSC UP state (**s**SPW-R: 85.9±1.4% and **c**SPW-R: 85.0±2.0%, mean±SEM, n=14 sessions in 4 mice^23^). **s**SPW-Rs concentrated near the end of the RSC UP state, which returned to a DOWN state immediately (<50 ms) after a large amplitude **s**SPW-Rs (**Figure 1F-H, J-L)**^23^. In contrast, the probability of occurrence of the first ripple of **c**SPW-R was lowest before the DOWN state (red line in **Figure 1L**). Following the DOWN-UP transition, ripples with the highest probability occurred ∼120 ms after UP state onset, and the subsequent ripples of **c**SPW-Rs occurred at similar intervals during the persisting UP state (**Figure 1L**), coupled with prolonged spiking activity of CA1 and CA3 pyramidal cells (**Figure 1F-H**), interneurons (**Figure S3**) and RSC neurons (**Figure 1F;** n=14 sessions in 4 mice).

We treated DOWN states shorter than 100 ms separately to avoid contamination from spindle-induced short silences (e.g., **Figure 1B**; **Figure S4C**). These short “DOWN” states corresponded to the peaks of the spindle waves (**Figure 1B**), and their troughs coincided with increased probability of SPW-Rs (**Figure 1L**). Together, these results suggest that **c**SPW-Rs preferentially occur during prolonged periods of cortical excitability and exhibit rhythmic structure, suggesting entrainment.

Consistent with this idea, **c**SPW-Rs were coupled to RSC spindle activity (8–12 Hz; **Figure 1D, I; Figure S4A-E**), co-occurring with spindles at significantly higher rates than **s**SPW-Rs (12.4±1.0% vs 5.0±0.3%, 2.5-fold, 1.6-fold increase, p=0.001, Wilcoxon signed-rank test, Cohen’s dz=1.92; 14 sessions in 4 mice). Ripple-containing spindles exhibited significantly higher peak power and were of longer duration than SPW-R-free spindles (**Figure S4F-I**). However, **c**SPW-Rs also occurred in the absence of spindles, suggesting a role for additional mechanisms, such as intrinsic resonance of hippocampal circuits^38^. To test this hypothesis, we optically stimulated CA1 parvalbumin interneurons from 2 to 40 Hz and revealed a strong resonance in the 5-12 Hz band for both putative pyramidal cells and interneurons (**Figure S5**). Thus, inter-ripple intervals may be paced by a combination of circuit resonance and phase-entrainment by cortical spindles.

### Clustered SPW-Rs enhance segregation of neocortical domains

Because **c**SPW-Rs were coupled to sustained RSC UP states and entrained by RSC spindles, we hypothesized that these clustered events represent a discrete unit of hippocampal output and therefore should exert coordinated, systems-level effects on additional downstream neocortical regions. To test this directly, we combined widefield imaging of the dorsal neocortex with silicon probe recordings in ipsilateral CA1, to examine the association of **c**SPW-Rs on global cortical activity and functional network structure. In Thy1-GCaMP6f mice (n=5), a 64- or 128-channel silicon probe was lowered through the left hemisphere into right hippocampal CA1, ipsilateral to a thinned-skull cranial window preparation (**Figure 2A)**^23^. Simultaneous optical and electrophysiological recordings were obtained during 1-2 hours of head-fixed sleep, and sleep quality in this condition was comparable to home-cage sleep^23^. Widefield recordings were hemodynamically corrected^39^ (see Methods) and aligned to the Allen Institute’s Common Coordinates Framework before parcellation into 27 regions of interest (**Figure S6)**^40^.

**Figure 2.**
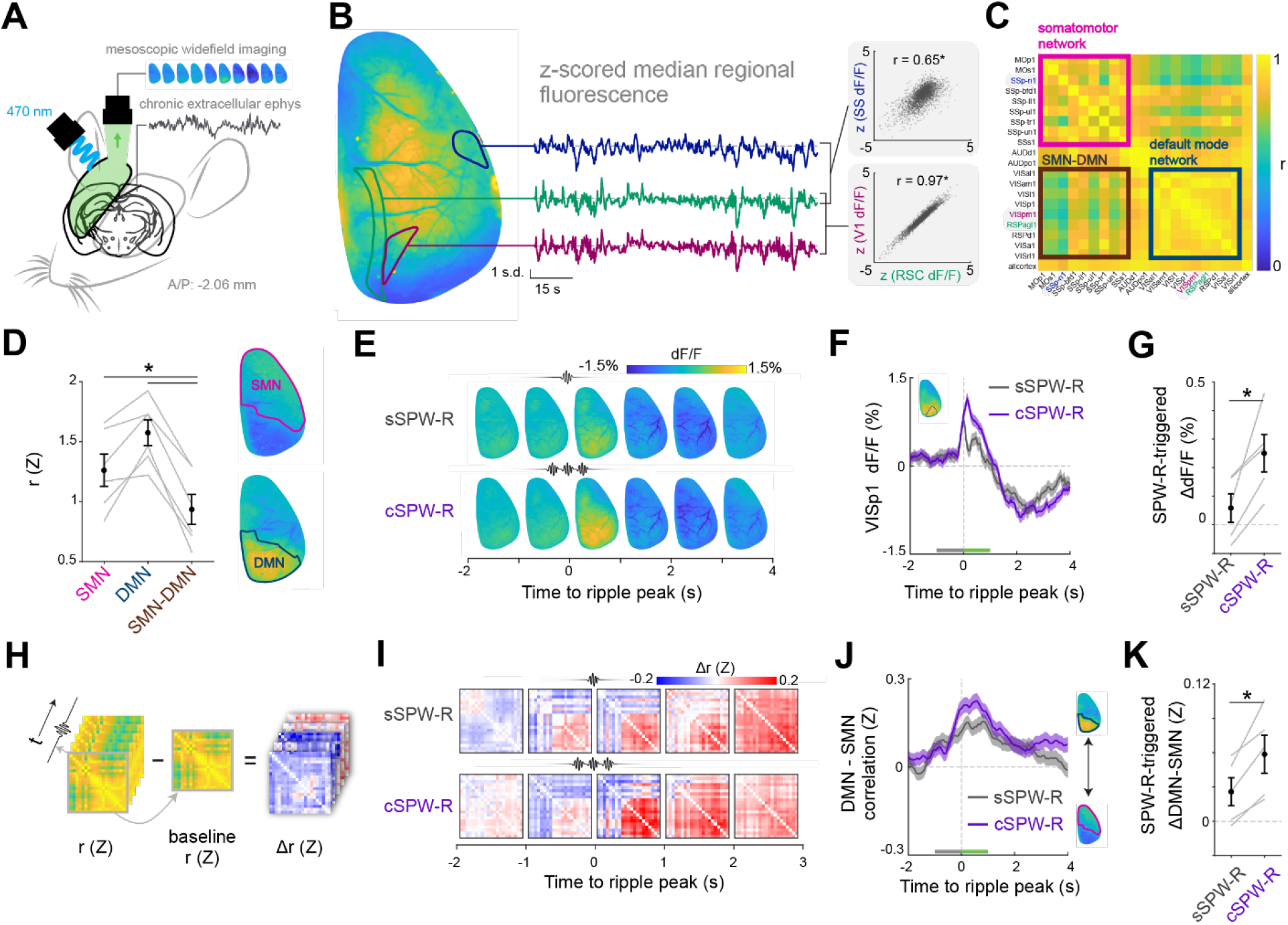
Differential engagement of global cortical activity by ripple clusters. **(A)** Schematic of recording setup. Mesoscopic widefield imaging (WF) of the right hemisphere of the dorsal cortical surface was performed in Thy1-GCaMP6f mice with simultaneous electrophysiological recordings in ipsilateral retrosplenial cortex and hippocampal CA1 using a silicon probe spanning both regions. **(B)** Example frame of WF recording (left). Median pixel intensity from three regions during NREM sleep (middle) and their significant pairwise correlations across all NREM periods of an entire sleep session (right). **(C)** Summary of NREM regional pairwise correlation for a single mouse. Regions from **B** are highlighted and canonical networks are labeled. **(D)** Summary of aggregate within- and between-network correlations (Z-transformed) across animals, showing that cortical subregions in different networks are less correlated than subregions in the same network (F_(2,10)_ = 47.27, p < 0.05; post-hoc comparisons of SMN and DMN vs SMN-DMN, t_(5)_ > 9.43*). Network boundaries are highlighted (right). **(E)** Sequence of frames depicting WF activity relative to peak power of **s**SPW-R or first **c**SPW-R in a cluster at t = 0, averaged across ripples (n_sSPW-R_ = 340 n_cSPW-R_ = 406) for a single animal. **(F)** SPW-R-triggered median fluorescence of a DMN representative subregion (VISp1) for the same animal. Inset depicts subregion outline. **(G)** Summary of the SPW-R-triggered change in fluorescence for VISp1 across animals (t_(4)_ = 3.70*). ΔdF/F was computed as the difference in mean fluorescence between the green and gray bars in **F**, corresponding to pre- and post-SPW-R periods across ripples for each mouse. **(H)** SPW-R-triggered changes in pairwise regional correlations (Z) were computed using a 1 second sliding window, followed by a baseline subtraction using the -5 to -2 second range prior to the ripple peak. **(I)** Sequence of matrices depicting SPW-R-triggered change in pairwise regional correlations, averaged across ripples for a single animal (same as in **E**). **(J-K)** SPW-R-triggered change in the balance between within-network DMN and SMN correlations (Z) for the same animal and **(K)** across animals (t_(4)_ = 4.57*). ΔDMN-SMN balance was computed as the difference in the mean traces between the green and gray bars in **J**, corresponding to pre- and post-SPW-R periods across ripples for each mouse. Shaded areas (**F**,**J**) and error bars represent SEM. *p < 0.05.

Pairwise correlations in regional fluorescence revealed two distinct cortical domains during NREM sleep, corresponding to the mouse default mode (DMN) and somatomotor networks (SMN)^41^, each exhibiting high within- and low cross-domain correlations (**Figure 2B-D**). The magnitude of fluorescent activity in DMN regions increased in the time window surrounding SPW-Rs^23^, and this effect was significantly enhanced by triggering cortical activity during **c**SPW-Rs, compared with **s**SPW-Rs (**Figure 2E-G**). Concurrently, somatomotor areas showed reduced or unchanged activity (**Figures S7-8A, B**). To examine short-timescale fluctuations in the functional organization of cortical networks, SPW-R-centered changes in pairwise regional correlations were computed using a 1-s sliding window (**Figure 2H**). Following **c**SPW-Rs, within-domain correlations for DMN and SMN regions diverged more than after **s**SPW-Rs (**Figure 2I, J; Figure S8C, D**), functionally segregating these networks and instating a DMN-dominant cortical mode (**Figure 2K**).

### Learning enhances cSPW-R-centered segregation of the default mode and somatomotor networks

Because SPW-Rs contribute to memory consolidation^4,18,27,42^, and **c**SPW-Rs are associated with transient segregation of cortical networks, we asked whether **c**SPW-Rs lead to stronger functional reorganization of DM-SM networks after learning. Mice (n = 3) were first trained to obtain water by running in a T-maze in which only one of two arms was accessible. After 14 days of training (‘*familiar*’), mice were tested in the same maze configuration for 50 trials before the second arm was made accessible (‘*novel*’) and the animal was rewarded for alternating between the left and right arms (**Figure 3A**). The mice learned the novel alternation task within the same session (**Figure 3B**) and continued to alternate with high proficiency the following day (**Figure 3C**) suggesting that an enduring memory of the task structure had formed after introducing the novel arm. To capture the effects of this learning on hippocampal-neocortical communication, we imaged the dorsal cortical surface and recorded from hippocampal CA1 during sleep before and after each behavioral session throughout training, as previously described (**Figure 2A**).

**Figure 3.**
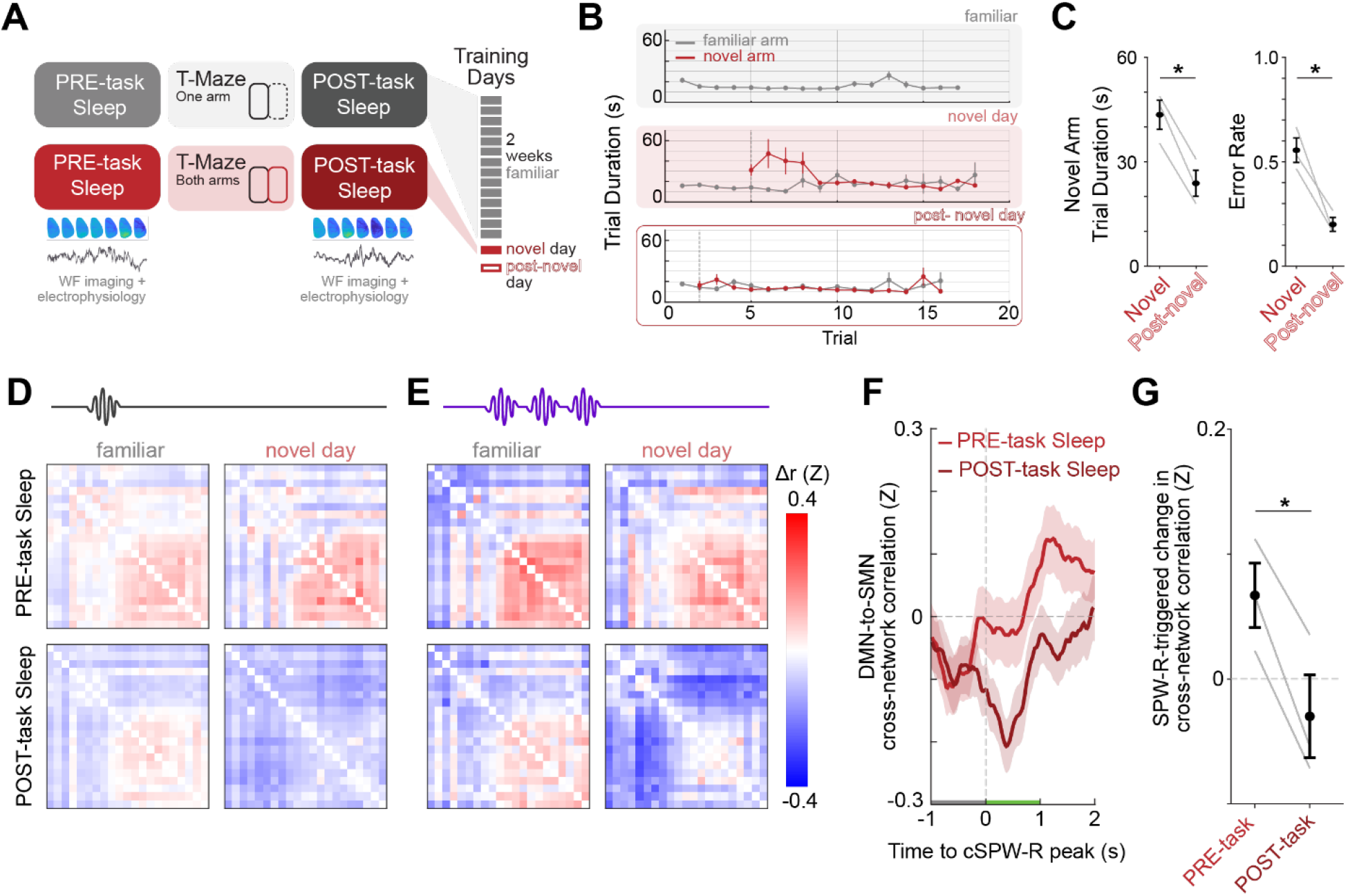
Ripple clusters isolate cortical networks during memory consolidation. **(A)** Schematic of behavioral training. Water-deprived mice were trained to run on one open arm of T-maze over two weeks (*familiar*), before the novel arm of the maze was introduced in a forced alternation phase of training (*novel/post-novel*). **(B)** Trial duration in 10-trial blocks across different training phases for a representative animal. **(C)** Summary of behavioral performance across animals. Early session novel arm trial duration (t_(2)_= -9.69*) and alternation error rate (t_(2)_ = -4.88*) decreased after a single exposure to the novel arm of the T-maze. **(D-E)** SPW-R-triggered change in pairwise regional correlations (Z) across behavioral conditions for a representative animal. Matrices represent Δr (Z) computed in the one-second window following ripple peak. **(F-G) c**SPW-R-triggered change in DMN-SMN between-network correlations before (*PRE-task Sleep*) or after (*POST-task Sleep*) a novel experience for the same animal and **(G)** across animals (t_(2)_ = - 7.61*). Change in cross-network correlation was computed as the -5 to -2 second baseline-subtracted aggregate between-network correlation. Summary statistics in **(G)** were computed as the difference in the mean traces between the green and gray bars in **F**, corresponding to pre- and post-SPW-R periods across ripples for each mouse. Shaded areas (**F**) and error bars represent SEM. *p < 0.05.

While **c**SPW-Rs remained associated with diverging within-network correlations (**Figure 2I-K; Figure 3D, E**), only post-learning **c**SPW-Rs were followed by a significant reduction in between-network DMN-SMN correlations (**Figure 3E-G; Figure S9**). This effect persisted even after controlling for changes in sleep depth before and after learning (**Figure S10**) and across alternative **c**SPW-R definitions (**Figure S11)**. In summary, **c**SPW-Rs are associated with learning-dependent functional isolation of key cortical networks. These data suggest that **c**SPW-Rs define transient windows during which hippocampal output may be broadcast with reduced cortical somatomotor interference, thereby creating conditions favorable for consolidation. Under this framework,clustered ripples should preferentially carry richer hippocampal representations than isolated events.

### sSPW-R and cSPW-Rs replay different aspects of waking-related activity

Because learning differentially modulated the cortical network effects of **c**SPW-Rs and **s**SPW-Rs, we hypothesized that these event types transmit distinct hippocampal content during consolidation. To test this, we compared the prevalence and spike structure of decoded reactivations across event types on high-density CA1 recordings from a third cohort of mice (18 sessions from 6 animals) performing a T-maze alternation task^43^. First, we used a hierarchical state space decoder^44^ to decode the 2D position of the animal (**Figure 4A**) during population synchrony events (i.e. a union of population burst and clustered ripple events) during NREM sleep. A shuffle test determined whether the co-activities of the neurons were well-described by the generative model using a spatial template (“on-manifold” events)^45^. We found that 14% of the SPW-Rs (n = 47877) were significantly better explained than by shuffled chance control. In PRE-task sleep and POST-task sleep, **c**SPW-Rs had a significantly higher fraction of on-manifold events compared to **s**SPW-Rs (**Figure 4B**; PRE: **s**SPW-Rs: 8%, **c**SPW-Rs: 9%; POST: **s**SPW-Rs: 12%, **c**SPW-Rs: 30%). The lengths of replayed trajectories (as measured by maximum displacement among continuous segments within a population synchrony window; **Figure 4C**) and the speed of replay correlated with the number of SPW-Rs contained in the synchronous events (**Figure S12**). This effect persisted even after controlling for duration of events, accounting for the large discrepancy between solo and cluster ripples (**Figure 4C**). The replayed content also differed depending on the maze zone. We distinguished replay of stationary locations, the reward areas and the delay area where animals paused without locomotion (‘within’; **Figure 4D**), from replay of the maze corridors traversed during ambulation (‘outside’; **Figure 4D**). Replays of stationary zones were more frequent during **s**SPW-Rs, whereas replays of the corridors were more frequent during **c**SPW-Rs, especially during POST-task sleep (**Figure 4E**).

**Figure 4.**
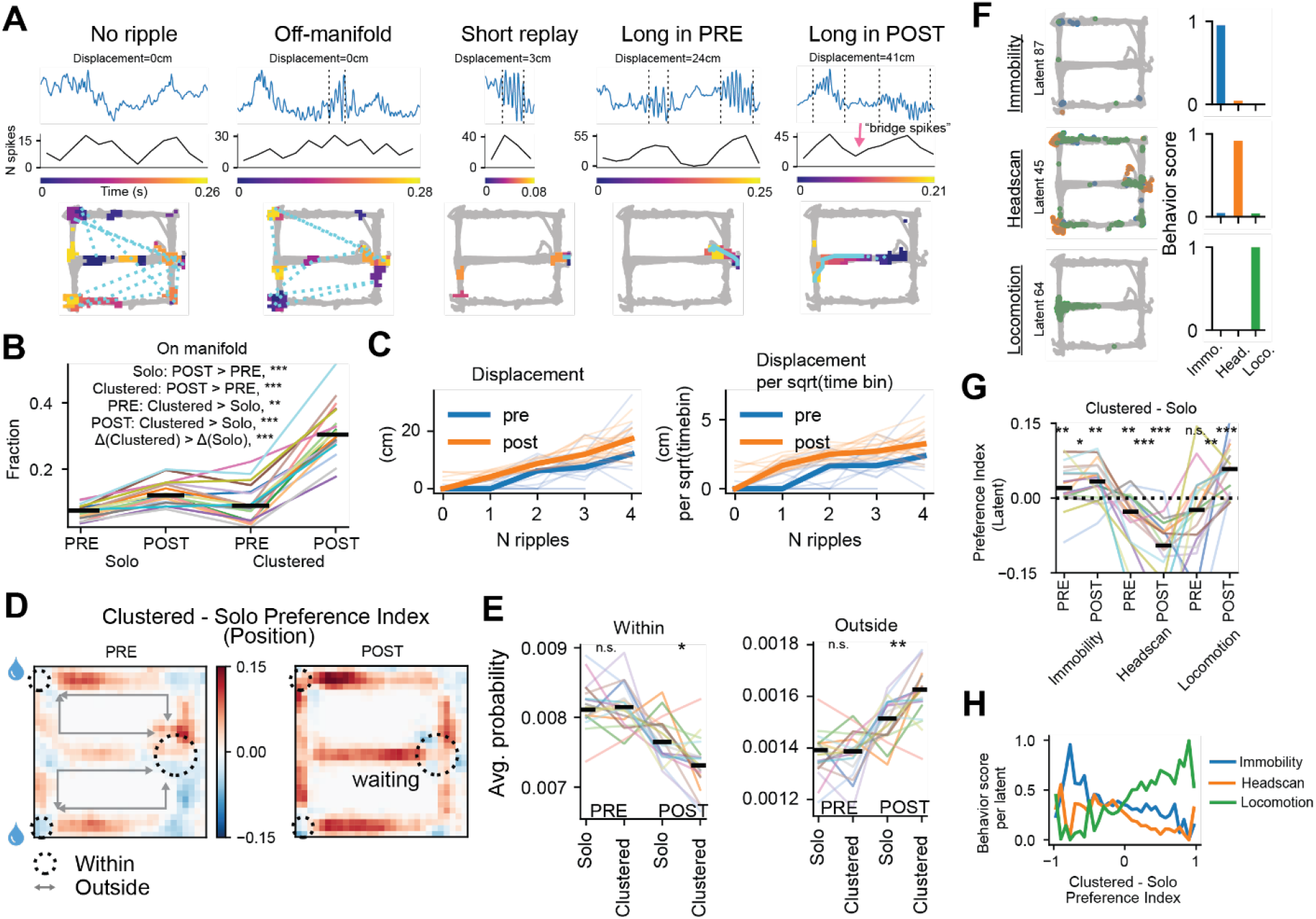
Replay differences between sSPW-Rs and cSPW-Rs. **(A-E)** Hidden Markov Model (HMM) decoding analyses. **(A)** Example events. LFP (top), population synchrony (middle) and events projected onto the maze (bottom). Colored patches are pixels with probability > 0.01 at each time bin, colored by time within the event. The cyan lines connect the maximum *a posteriori* (MAP) pixels across time. Solid lines, continuous segments; dotted lines, fragmented segments. Note long, continuous trajectory replays in the last two columns. (**B**) The fraction of on-manifold events within each sub-category of events. On-manifold is defined as passing the shuffle tests, which assess whether the spatial tuning curves accurately describe the population co-activity during synchronous events. Each colored line is one session. Black bars are medians across sessions. **c**SPW-Rs have a significantly higher fraction of on-manifold events than **s**SPW-Rs during both PRE-task and POST-task sleep. The gain is also significantly higher during POST-task sleep. (**C**) Metrics of replayed trajectories as a function of the number of SPW-Rs contained in a synchronous event: displacement (left) and displacement divided by the square root of the number of time bins (right), separated into PRE-task (blue) and POST-task (orange) sleep. Thin lines, individual sessions; thick lines, median across sessions. (**D**) Heatmaps of Preference Index at each position bin. Probabilities during **c**SPW-Rs and **s**SPW-Rs were first averaged over time for a given epoch; PRE-task sleep (left) and POST-task sleep (right). Preference Index was computed as (c-s)/(c+s). The maps were then aligned and averaged across sessions. Reward ports are marked with water droplets. Black dotted circles indicate regions with low locomotor speed at reward ports and in the delay area. (**E**) Spatio-temporal average of the probability within (left) or outside (right) of the circled low-locomotion regions in D, separated into event epochs (PRE or POST) and types (c or s). Each line is one session. Black bars, medians across sessions. **c**SPW-Rs decode events with significantly higher probability during locomotion in corridors (i.e., outside) during POST-task sleep. (**F-H**) JumpLVM analysis results. (**F**) Left: example activation patterns of different types of latent bins. Each dot marks the 2D position of the animal when that latent bin achieves MAP, colored by the behavior type at that time (blue: Immobility, orange: Headscan, green: Locomotion). Right: behavior score of that latent bin. Even though the headscan latent (middle row) still activates during locomotion occasionally, it occurs much more reliably during headscans (i.e. occuring in most headscan events, but only sparsely during locomotion), which is captured by the behavior score. (**G**) Clustered-Solo Preference Index per latent bin, separated into PRE- and POST-task sleeps and latent types. Each colored line is a session; solid black short lines, medians across sessions. Dotted line, no preference for **c**SPW-Rs or **s**SPW-Rs. Headscan latents occur primarily during **s**SPW-Rs, while Locomotion latents gain preference for **c**SPW-Rs. (**H**) Behavior scores for each behavior type (color) per latent bin as a function of the c-s Preference Index, aligned and averaged across all sessions.

Even at the same maze position, the hippocampal neural representation depended on the animal’s behavior, suggesting that a purely spatial decoder conflates behaviorally distinct population patterns that happen to share the same location^45^. To complement the trajectory-based replay analysis, we asked whether the reactivated population patterns (independent of their spatial content) also differed between **c**SPW-Rs and **s**SPW-Rs. We thus used an unsupervised method, the Jump Latent Variable Model (JumpLVM^45^; **Figure S13A-B**), to extract latent variables (i.e. patterns of population activities) during the task-epoch. We then characterized the latent by behavior, decoded the latent during synchronous population events, and performed a similar shuffle test as in the supervised case to select “on-manifold” events for further analysis. The JumpLVM reduces population activities to a 1-D nonlinear latent manifold (discretized into bins) without requiring the latent to correspond to external labels, such as position. Behavioral correlates (templates) are learned from the correlational structure of population activities and the smoothness condition, quantified by the similarity of population firing between adjacent latent bins. Based on how frequently each latent bin activates during each behavior type (during the task epoch), we scored and classified each latent bin into Immobility, Headscan, and Locomotion (**Figure 4F**; **Figures S3, 13C**). Headscan latents preferred **s**SPW-Rs during both PRE- and POST-task sleep, while Locomotion latents preferred **c**SPW-Rs, but only during POST-task sleep (**Figure 4G, H**; **Figures S13D, 14**). On average, Immobility latents preferred **c**SPW-Rs (**Figure 4G**), but there was a large spread (**Figure 4H**). Together, our results highlight differences in replay content between **c**SPW-Rs and **s**SPW-Rs: **c**SPW-Rs are associated with replaying long trajectories during locomotion, whereas **s**SPW-Rs are more strongly associated with activity patterns during non-locomotion behaviors. The latent reactivation analysis further corroborates the conclusion that the difference in replay content between **c**SPW-Rs and **s**SPW-Rs cannot be explained by the difference in event durations.

## Discussion

Our findings, together with prior work, support a coordinated cascade linking cortical slow oscillations, thalamocortical spindles, and hippocampal SPW-Rs during sleep. A fraction of cortical DOWN-UP transitions occur as highly synchronous K-complexes^46–49^, which can trigger thalamocortical spindles^50–52^. The synchronous spiking associated with these spindles can, in turn, entrain clustered SPW-Rs that are phase-locked to spindle troughs (**Figure 1D**)^3,5,23,53,54^. During the UP state, sustained neocortical activity and SPW-R clustering enable reverberating hippocampal-cortical spike exchange^3,55,56^; providing a window for structured information transfer and the sequential ordering of cortical representations^57^. This communication window is terminated when the final ripple of a cluster, or a large-amplitude **s**SPW-R drives the network back into a DOWN state (**Figure 1K, L**)^23,58^. Following learning, these clustered events are accompanied by enhanced segregation of cortical default mode and somatomotor networks (DMN and SMN) (**Figure 3)**, and by longer, more continuous replay trajectories (**Figure 4**)^20,28^, consistent with the concatenation of waking experience into coherent episodic sequences^20,59,60^.

The ripple cluster hypothesis builds on prior work and unifies several observations. **c**SPW-Rs transiently enhance the functional segregation of SMN and DMN, effectively deepening an offline cortical state in which associative regions such as the hippocampus and DMN are privileged while somatomotor areas are disengaged. This pattern mirrors the task-negative organization of the DMN^61^ and suggests that ripple clusters create protected windows for hippocampal-neocortical dialogue, minimizing interference from ongoing sensory processing^62,63^. For example, presenting sounds during sleep biases subsequent hippocampal replay content but simultaneously suppresses overall ripple incidence^56^, while minor local body movements during NREM microarousals can robustly suppress SPW-R rates^64^. These data suggest that both intrinsically and extrinsically driven cortical activity can influence ongoing hippocampal-DMN dynamics, necessitating a mechanism that biases cortical dynamics away from bottom-up interference, particularly during early consolidation following novel experiences. From a complementary learning systems perspective,transient network segregation may also protect against catastrophic interference by restricting when and where hippocampal information is broadcast to the cortex. By limiting cross-network coupling during early consolidation, **c**SPW-Rs could enable the gradual integration of new traces without disrupting established cortical representations^54,65,66^. SPW-R-coupled DOWN states may provide a protected period for cortical consolidation, preventing interference from subsequent hippocampal inputs while synaptic modifications induced by the preceding cluster are stabilized. Thus, DOWN states may help in segregating distinct memory traces during consolidation.

Several lines of convergent mechanistic evidence suggest that slow oscillations and thalamocortical spindles actively entrain hippocampal ripples, thereby promoting memory consolidation. In humans and other mammals, K-complexes can be triggered by sensory stimulation during NREM sleep at modality-specific sites^52^, and auditory closed-loop entrainment of slow oscillations enhances memory (“targeted memory reactivation”;^9,12,67^, but see Henin et. al. 2019^68^). Similarly, artificial coupling of SPW-Rs to slow oscillations strengthens cortical responses to ripples^17^, while optogenetic induction of spindles during the UP state reliably triggers hippocampal SPW-Rs^69^, with both manipulations enhancing hippocampus-dependent memory consolidation. Consistent with this role, spindles structure the temporal organization of SPW-Rs, as ripples are phase-locked to spindle troughs (**Figure 1D**)^2,3,5,11,70^ and contribute to a prominent peak in the ripple-autocorrelogram, superimposed on an otherwise stochastic distribution (**Figure 1C**)^20^. Intrinsic resonant properties of hippocampal circuits may further amplify this entrainment (**Figure S5**), contributing to the increased amplitudes observed within **c**SPW-Rs (**Figure S1C**) and to the stronger association between large-amplitude SPW-Rs and successful consolidation^71^. Finally, this spindle-ripple coupling is likely mediated through the entorhinal cortex, since UP states and spindles entrain SPW-Rs^53,72^, and layer 3 inactivation reduces SPW-R incidence^20^.

**c**SPW-Rs may support the binding of sequential experiences into extended episodes, as the intervals between clustered ripples fall within the time window of NMDA receptor-dependent plasticity^73^. This temporal structure could allow consecutive events to be linked and associations to form across ripples. Beyond simple sequencing, **c**SPW-Rs may also integrate new memories into existing knowledge^60^ through hippocampal-neocortical reverberation during extended UP states. Consistent with this integrative role, behavioral replay is differentially organized across ripple types. Locomotor trajectories preferentially occur during **c**SPW-Rs and increase after learning, whereas head-scanning and immobility sequences are primarily expressed during **s**SPW-Rs, and dominate before learning. While debates about ‘preplay’ and ‘replay’ have focused on awake ripples and prospective vs. retrospective coding^30,34,74^, our findings reveal analogous heterogeneity within sleep-associated SPW-Rs. Just as awake ripples show functional diversity linked to behavioral demands, sleep ripples segregate into distinct subtypes (**c**SPW-Rs and **s**SPW-Rs) that engage cortical networks differentially and may consolidate different behavioral experiences (**Figure 4**). This suggests that state-dependent specialization (wake vs. sleep) and pattern-dependent specialization (clustered vs. solo) operate as complementary organizing mechanisms for hippocampal-neocortical communication. Overall, these findings indicate that ripple clustering organizes hippocampal output into temporally extended, sequential structures through coordinated interactions with thalamocortical spindles, providing a substrate for compositional computation and abstraction^57,75^.

## Data availability

All data are available in the manuscript or the supplementary materials or are publicly available in the Buzsaki Lab Databank, https://buzsakilab.com/wp/database/.

## Code availability

All code used for data pre-processing is available at https://github.com/buzsakilab/buzcode. The code for analyzing neural data, visualization, and application of the supervised neuronal classifier will be available at Zenodo after manuscript acceptance.

## Acknowledgements

We thank Joaquin Gonzalez, Gergely Komlósi, JingJing Liu, Heechul Jun and members of our laboratory for helpful comments on the project.

## Author contributions

Conceptualization: M.V., G.B.; Funding acquisition: G.B., E.Y., C.L., N.P.; Investigation: M.V., C.L., Z.Z., N.P., E.C., K.M., D.A. Project administration: M.V., G.B.; Supervision: M.V., C.L., G.B.; Visualization: M.V., C.L., Z.Z. Writing – original draft: G.B., M.V.; Writing – review & editing: G.B., M.V., C.L., Z.Z and all other authors.

