## Supplementary File for "Sharp wave-ripple clusters enhance hippocampal-neocortical engagement for memory consolidation"

**The PDF file includes:**

Materials and Methods

Figs. S1 to S15

Supplementary Tables 1 to 3

### Materials and Methods

#### In vivo animal experiments

All experiments were approved by the Institutional Animal Care and Use Committee at New York University Medical Center. Mice and rats were housed in a vivarium on a 12-hour light/dark cycle, four per cage before surgery and individually after surgery.

#### Surgery for RSC, CA1, CA3 recordings

For details on animal surgery (adult, male n = 3 and female n = 1; C57Bl6), training, recording, data preprocessing, we refer to Gonzalez et al. (2026)[76]. In short, a ground screw was placed above the cerebellum and a 4-shank Neurpixels2.0[37] probe was implanted at 2 mm posterior to the bregma and 1.5 mm lateral to the midline on the left side. The probe was lowered 6 mm from the brain surface at a 60 degrees angle, targeting the RSC, CA1, and CA3[23,76]. The microdrive was secured to the skull[77]. The craniotomy was sealed with a layer of Duragel (Cambridge Neurotech). We used SpikeGLX (https://billkarsh.github.io/SpikeGLX/) to optimize channel selection, maximizing the number of well-isolated single units in each target region. The collected data were digitized at 30 kS s−1 using SpikeGLX software and a PXI acquisition system.

#### Surgery for optogenetic recordings

Transgenic mice (adult males, n = 2) expressing channelrhodopsin-2 selectively in parvalbumin-expressing inhibitory cells were implanted with a 3D-printed headpost[78] for awake, head-fixed experiments. Animals were allowed to walk freely on a low-friction rodent-driven treadmill during recording sessions[79]. A 64-channel silicon probe (H2, Cambridge Neurotech) attached to a 200-μm optic fiber[80] was used to record neuronal activity. The day before recording, a craniotomy was performed (2 mm posterior to Bregma and 1.5 mm lateral to midline) and the dura was removed. After surgery, the craniotomy was sealed with Kwik-Sil (World Precision Instruments) until recording. On the day of recording, the animal was head-fixed, the craniotomy was cleaned, and the headpost was filled with sterile saline. The ground of the probe was connected to the header pin, and the probe was inserted to the target depth using a manual micromanipulator (MM-33, Sutter Instruments). Electrophysiological signals were continuously monitored during insertion. The collected data were digitized at 20 kS $s^{-1}$ using an RHD2000 recording system (Intan technologies). We waited at least 15 min after reaching the target depth. Baseline sessions (30 min) and optogenetic stimulation sessions were recorded from each mouse.

Light delivery was controlled using a custom open-source hardware driver. A Teensy 3.2 USB development board was used with an Open Ephys Cyclops LED driver board. The Cyclops board was configured to drive a laser diode (L450G3, Thorlabs). To characterize frequency-dependent responses (resonance) while avoiding history-dependent effects often associated with continuous chirp signals, we used a randomized "pure frequency" stimulation protocol. Stimulation waveforms consisted of sinusoidal intensity modulations at discrete frequencies. Frequencies were defined in a pre-generated session header file to ensure reproducibility. Frequencies ranged from 1 to 40 Hz. The trial structure was as follows: (1) Stimulation Epoch: 1 second of sinusoidal stimulation at a specific target frequency, (2) Inter-Trial Interval: 2 seconds of no light output between trials to allow neural activity to return to baseline, (3) Order: frequencies were presented in a pseudo-randomized order to prevent adaptation to a predictable sequence. The stimulation waveforms were computed in real time on the Teensy microcontroller using custom firmware. The firmware used the *micros()* clock function to ensure high temporal precision. For each defined frequency, the instantaneous current intensity was calculated using a sinusoidal function:

$$I(t) = A\left(\frac{1+\sin(2\pi f t)}{2}\right),$$

where $A$ represents the defined amplitude scaling factor. This scaling factor was adjusted for each animal based on evoked responses with square pulses before the baseline recordings.

**Optogenetic tagging of PV cells**

To optogenetically tag PV cells in the hippocampus, PV-Cre::Ai32 mice were used. 2 ms light pulses were delivered every 200 ms for at least 5000 times. Sharp onsets and offsets were associated with a photoelectric artifact. To prevent such artifacts from propagating to spike sorting and unit identification, raw data were clipped out in the interval shortly before the onset (0.15 ms) and after the offset (0.8 ms) of each brief pulse[43]. To identify ChR2-expressing neurons, we tested recorded neurons for both characteristic firing latencies and reliable poststimulus responses. Latency effects were quantified using SALT[81], comparing the distribution of latencies to first spike in 10-ms windows after each pulse against baseline latency distributions (n=200) from randomly selected timepoints before stimulation ($P \leq 0.001$ considered significant). To test firing reliability while controlling for polysynaptic effects, we computed peristimulus time histograms (1-ms bins) smoothed with a hollowed Gaussian (15 ms s.d.) and assessed whether spike counts in 3 bins surrounding peak poststimulus firing significantly exceeded both baseline counts (Poisson distribution) and pre-stimulus firing at matched lags ($\alpha=0.001$, Bonferroni corrected).

**Surgery for dorsal, intermediate and ventral CA1 recordings**

To record from multiple locations along the longitudinal axis simultaneously, one or two Neuropixels2.0 electrodes were chronically implanted in rats (male n=1, female n=2, Long Evans, 300-350g) in three configurations. For dorsal-intermediate recording, the probe targeting dorsal CA1 was implanted 3.4 mm posterior to Bregma and 2.75 mm lateral to midline at a 15-degree angle. The other probe targeting intermediate CA1 was implanted 5.2 mm posterior to Bregma and 3 mm lateral to midline at a 15-degree angle. For one dorsal-ventral recording, one probe was angled 25 degrees rostral from vertical at 4.54 mm posterior to Bregma and 2.48 mm lateral to midline, passing through dorsal and ventral CA1. In the last configuration, the probe targeting dorsal CA1 was implanted 2.75 mm posterior to Bregma and 2.25 mm lateral to midline. The other probe targeting ventral CA1 was implanted 5.2 mm posterior to Bregma and 5 mm lateral to midline. A stainless-steel screw was implanted above the cerebellum as ground/reference. The probes were protected by a 3D-printed cap[77]. Animals were recorded in their homecage. The collected data were digitized at 30 kS $s^{-1}$ using SpikeGLX software and a PXI acquisition system.

**Surgery for widefield recordings**

For details on animal surgery (adult males, n=5), data preprocessing, and spike sorting, see Swanson et al. (2025)[23]. In brief, skull thinning was performed over the entire dorsal right hemisphere, from the frontal cortex to the posterior visual cortex. The skull was coated with a thin layer of cyanoacrylate (gel-type Loctite Super Glue) to index-match the rough surface of the skull, prevent bone regrowth, and provide mechanical protection. A thin coating of clear nail polish was then applied on top of the cyanoacrylate to further index-match the skull's surface and reduce scatter. Finally, a ground screw coupled with a 0.005" stainless steel wire (A- M Systems, #792800) was implanted in the skull above the cerebellum, and a custom 3D-printed head post was fixed to the surface of the skull using dental cement. In a second surgery, performed 5 days to 1 week after the first, mice were chronically implanted with a 128-channel multi-shank silicon probe (H3 or H20; Cambridge Neurotech) at a 60-degree angle, mounted on a microdrive [77]. The probe spanned area CA1 of the hippocampus and retrosplenial cortex ipsilateral to the imaging field of view, after being lowered from a craniotomy at -2.06 mm A/P -1.06 mm M/L, and -2.13 mm D/V relative to bregma in the contralateral (left) hemisphere. The probe and microdrive were then enclosed within the remaining components of the 3D-printed head post.

**Widefield recordings**

The following section is adapted from Swanson et al. 2025[23] (**Figure S6**). Widefield imaging was performed on the right hemisphere of head-fixed mice through a thinned-skull preparation using the MVX-10 Macroscope (Olympus) during simultaneous chronic electrophysiological recordings from the ipsilateral retrosplenial cortex (RSC) and hippocampal CA1 region. Data were acquired with an Andor Zyla sCMOS camera using dual-wavelength LED illumination (cool LED PE-4000 system) and controlled via NIS Elements software (Nikon). 16-bit images were acquired at 66.66 Hz in global-shutter mode. Illumination alternated between 470 nm and 525 nm per frame, and emitted light was collected through a 500-nm long-pass emission filter.

For more details about the denoising, see Peters et al. 2021[82]. All wide-field data were de-noised via compression using singular value decomposition of the form F = USVT. The input to the SVD algorithm was F, the pixels x time matrix of fluorescence values. The outputs were U, the pixels x components matrix of template images; V, the time x components matrix of component time courses; and S, the diagonal matrix of singular values and the top 20 components were retained. Hemodynamic effects on fluorescence were removed by regressing out a green 525 nm widefield channel, reflecting changes in total blood volume, from the calcium-dependent signal obtained with blue illumination (470 nm). To achieve this, both signals were band-pass filtered in the range 7–13 Hz (heartbeat frequency, expected to have the largest hemodynamic effect). Pixel traces for blue illumination (470 nm) were then temporally resampled to be concurrent with green illumination (525 nm, as colors were alternated), and a scaling factor for the given pixel-wise regression was fit across colors for each pixel. The scaled fluorescence traces from 525 nm green light illumination were then subtracted from the fluorescence traces emitted from 470 nm blue light illumination. To correct for slow drift, hemodynamic-corrected fluorescence was then highpass filtered over 0.01 Hz and ΔF/F0-normalized by dividing by the average fluorescence at each pixel within the session. Wide-field videos were then aligned across sessions by rigid registration of the average hemodynamically corrected image to the Allen Common Coordinate Framework [40] using four manually selected anatomical landmarks (marked during surgery).

Averaging and statistical comparisons of pairwise-regional correlations were performed on Z-transformed Pearson's r scores to approximate normality, stabilize variance and ensure valid parametric comparisons.

**Sharp wave-ripple detection**

A single LFP signal was bandpass filtered in the ripple band (130-250 Hz), and the normalized squared signal was calculated. SPW-R peaks were detected by thresholding the normalized squared signal at 5x SDs above the mean, and the surrounding SPW-R start and end times were identified as crossings of 2x SDs around this peak. Ripple events lasting >200 ms were excluded to minimize artifacts. The detected events were further manually inspected using NeuroScope2[83], and noise artifacts and false positives were removed, typically arising from electrical or movement artifacts. A reference channel outside the pyramidal layer of CA1 was also used to filter out false-positive ripple events. The channel used for ripple detection was visually defined based on individual ripples, where ripple power was highest.

**Brain state scoring**

Brain state scoring was performed as described by Watson et al. (2016)[84]. In brief, spectrograms were constructed using a 1-s sliding window and a 10-s fast Fourier transform of the LFP at log-spaced frequencies between 1 Hz and 100 Hz. Three types of signals were used to score states: broadband LFP, narrowband theta-frequency LFP, and electromyogram (EMG). For the broadband LFP, principal component analysis was applied to the Z-transformed spectrogram (1–100 Hz). In all cases, the first principal component was based on power in the low-frequency band (<32 Hz). Theta dominance was defined as the ratio of power in the 5–10 Hz and 2–16 Hz bands in the spectrogram. EMG was extracted from the intracranially recorded signals by computing zero-lag correlation coefficients (r) between 300 and 600 Hz filtered signals (using a Butterworth filter at 300–600 Hz with filter shoulders spanning 275–625 Hz) recorded at all sites. Next, all states were inspected and curated manually, and corrections were made when discrepancies between automated scoring and user assessment occurred.

**Spindle detection**

Sleep spindles were identified from local field potentials (LFPs) recorded in the retrosplenial cortex (RSC) using a wavelet-based approach adapted from Johnson et al. (2012) and Sullivan et al. (2014)[85,86]. This procedure estimates instantaneous power in the spindle frequency range (5–15 Hz) and detects epochs of sustained rhythmic activity during non-rapid eye movement (NREM) sleep. LFPs were downsampled to 1250 Hz and filtered between 5 Hz and 15 Hz using a zero-phase finite-impulse-response (FIR) band-pass filter. This frequency range was chosen because sleep spindles in mice occur at lower frequencies[87] than the canonical human spindle band (11–16

Hz); in our data, the population mean instantaneous spindle frequency was 9.53 ± 2.33 Hz (n = 14 sessions; **Figures S4D**). Sleep state annotations were imported from SleepState files, and only NREM segments were considered for detection. For each LFP trace, a complex Morlet wavelet transform (center frequency ≈ 11 Hz, wavelet parameter = 6) was computed over scales corresponding to 5-15 Hz. Wavelet power was averaged within NREM epochs to obtain the mean and standard deviation across scales. The instantaneous spindle-band power was then z-score normalized relative to this NREM baseline. Candidate spindle epochs were defined as contiguous intervals in which the z-scored spindle power exceeded 1.4 SD for ≥ 350 ms. Adjacent events separated by < 100 ms were merged. Events with peak z-power < 2 SD were discarded. To ensure rhythmic integrity, individual oscillatory cycles were examined within each candidate epoch; spindles containing any cycle longer than 125 ms (≤ 8 Hz) were excluded. For each spindle, onset, offset, and peak times were extracted from the z-scored envelope; corresponding trough times were determined from the filtered LFP waveform.

**Inter-ripple interval analysis** and **SPW-R cluster detection**

SPW-Rs were detected using the bz_FindRipples algorithm (Buzsáki Lab GitHub). SPW-R clusters were defined as chains of ripple events separated by short inter-event intervals (IEIs). To determine an objective, physiologically valid threshold for clustered SPW-R (cSPW-R) detection, we performed a rigorous statistical analysis of the pooled IEI distribution across all sessions (n = 25 sessions, n = 7 mice, total ripples = 86354). Timestamps of detected SPW-Rs were sorted, and IEIs were calculated as the time difference between the peaks of consecutive ripples. We fitted the pooled IEI distribution in logarithmic space with a double Gaussian model to account for its log-normal nature. The distribution was binned into 100 logarithmically spaced bins, and normalized histogram counts were fitted with the following model:

$$f(x) = a_1 exp\left(-\frac{(x - \mu_1)^2}{2\sigma_1^2}\right) + a_2 \exp\left(-\frac{(x - \mu_2)^2}{2\sigma_2^2}\right)$$

where $x$ represents $\log_{10}$-transformed IEI values, $\mu_1$ and $\mu_2$ are the centers of the two Gaussian components, $\sigma_1$ and $\sigma_2$ are their standard deviations in log-space, and $a_1$ and $a_2$ are their respective amplitudes. Parameters were estimated using nonlinear least-squares optimization with bounded constraints: amplitudes [0, 2], centers [$\log_{10}(0.01)$, $\log_{10}(100)$] seconds, and standard deviations [0.01, 2] in log-space. The first Gaussian component (fast component) captured the cluster regime, representing rapid sequences of SPW-Rs, while the second component (slow component) captured the background regime of isolated events. The crossing point between the two Gaussians (where their probability densities were equal) was calculated as the intersection of the fitted curves and used as a natural boundary between clustered and isolated events. Clustered SPW-Rs (**c**SPW-Rs) were defined by an IEI of <180 ms, a threshold that closely aligns with the upper intersection point (177 ms) where the cluster (G1) and solo (G2) populations are equiprobable. At 180 ms, the cluster component amplitude remains at 57.4% of its peak, and the threshold captures 76.1% of the total area of the cluster distribution, ensuring high sensitivity for clustered events. For solo SPW-Rs (**s**SPW-Rs), we set a conservative threshold of ≥500 ms. This window extends beyond the 1% decay point of the cluster distribution (343 ms), at which point the cluster component amplitude is negligible (0.014% of peak).

**Sleep depth and spectral analysis**

To characterize the local network state during SPW-R events, we analyzed the Local Field Potential (LFP) from the channel with the highest ripple amplitude (i.e., the middle of the CA1 pyramidal layer). The raw LFP was band-pass filtered into Delta (1–4 Hz) and Theta (6–9 Hz) frequency bands using a 3rd-order Butterworth filter. The instantaneous amplitude of each band was extracted via the Hilbert transform. A continuous Theta/Delta ratio signal was calculated by dividing the instantaneous Theta amplitude by the Delta amplitude. The resulting ratio trace was downsampled to 50 Hz for analysis. To compare event-related dynamics, the Theta/Delta ratio was extracted around each ripple event with a window of ±2 seconds. For Solo ripples, the window was aligned to the timestamp of the ripple peak power. For Cluster events, the window was aligned to the temporal center of the cluster duration (defined as $(T_{start} + T_{end})/2$). Event-triggered averages were calculated for each session, and statistical comparisons were performed using a paired Wilcoxon signed-rank test on the mean ratio values at t=0.

**Single unit analysis**

A concatenated signal file was prepared by merging all recordings from a single animal on a single day. Putative single units were first sorted using Kilosort2.5[88] and then manually curated using Phy (https://phy.readthedocs.io/en/latest/).

**Cell type classification**

In the processing pipeline, cells were classified into two putative cell types: interneurons and pyramidal cells. Interneurons were identified using two criteria. We labeled single units as interneurons if their trough-to-peak latency was <0.425 ms, or if it was >0.425 ms and the rise time of the autocorrelation histogram was >6 ms. The remaining cells were assigned as pyramidal cells. Autocorrelation histograms were fitted with a triple-exponential equation to supplement the classical, waveform feature-based single-unit classification (https://cellexplorer.org/pipeline/cell-type-classification/)[89]. Spike bursts were defined as groups of spikes with interspike intervals < 9 ms.

**Behavioral training and experimental design**

**Widefield imaging dataset**

Mice were habituated for ≥1 week to head-fixation using a modified rivet system[78], with fixation duration gradually increasing from 10 min to 2 h over days. Animals readily slept under head-fixation, confirmed by electrophysiological markers and spectral measures of sleep depth, with sleep quality comparable to home-cage conditions[23] (**Figures S7,10**). In parallel, mice were tethered and acclimated to a T-maze. During habituation, only one arm was accessible while the other remained blocked. By the end of habituation, daily sessions consisted of morning pre-sleep head-fixation (1–2 h), maze exposure (1 h), and afternoon post-sleep head-fixation (1–2 h).

Under water restriction, mice were then trained to run the maze for reward. Trials began in the home area of the maze. After a brief delay, barriers opened to allow the mouse to run along the central arm and turn at the T-junction, where one arm was accessible and the other was blocked. After reward delivery, a rear barrier prevented reversal and the home door reopened to allow initiation of the next trial. Training continued until animals reliably obtained ≥150 rewards per hour.

After reaching criterion performance, experimental sessions were performed. In the 'familiar' condition, electrophysiology and widefield calcium imaging were recorded before and after behavior, with electrophysiology recorded throughout task performance. In the novel condition, these recordings were repeated but the previously blocked arm was introduced after 50 trials, requiring forced alternation to obtain reward. The novel arm differed from the familiar arm in floor texture and in visual markers along the maze wall. In post-novel sessions, the second arm was introduced after only 10 trials. Behavioral performance was quantified using arm-specific trial duration and forced-alternation error rates.

This behavioral paradigm was designed to induce spatial learning through modification of an established cognitive map rather than exploration of an entirely novel environment. Structural changes in maze geometry, including barrier insertion or removal, are known to reorganize hippocampal spatial representations as newly accessible regions are incorporated into the cognitive map [90–92]. By introducing a previously inaccessible arm after animals had formed a stable representation of the familiar trajectory, this paradigm provided a controlled means to examine neural activity associated with updating an existing spatial framework. Because learning occurred relative to a familiar baseline, post-learning activity could be compared against a well-defined reference state, isolating neural dynamics associated with updating established representations. We therefore focused on neural activity during post-learning sleep following exposure to the novel arm to assess how hippocampal output related to spatial updating influences large-scale cortical activity.

**Dataset and task, related to Figure 4.**

For details on animal surgery, training, recording, data preprocessing, and spike sorting/cell body segmentation, we refer to Huszar et al. (2022)[43]. In brief, we used chronic silicon probe recordings from the hippocampal CA1 region. Animals were trained on a spatial alternation task in a figure-eight maze. Animals were restricted to water before the start of experiments and familiarized with a customized 79x79 $cm^2$ T-maze raised 61 cm above the ground. Over several days after the start of water deprivation, animals were shaped to visit alternate arms between trials to receive a water reward. A 5-s delay in the start area (delay area) was introduced between trials. The position of head-mounted red LEDs (light-emitting diodes) was tracked with an overhead camera at a frame rate of 30 Hz. Animals were required to run at least ten trials along each arm (at least twenty trials total) within each session. In all sessions that included maze behavior, animals spent 120 min in the homecage before running on the maze and another 120 min in the homecage afterward for sleep recordings. All behavioral sessions were performed in the mornings (start of the dark cycle). We used 18 sessions with the highest cell counts (putative pyramidal cells ranging from 132 to 422). Sharp wave ripple (SPW-R) and sleep state scoring are provided in the dataset (Huszar, 2022)[43].

**Behavior classification**

2D velocity (vx and vy) was computed via finite difference from 2D position tracking and then smoothed with a Gaussian filter (standard deviation 300 ms). Speed was computed as:

$$v = \sqrt{v_x^2 + v_y^2}$$

Immobility was defined as periods when speed was below 2.5 cm/s. Headscan was defined by the following procedure (adapted from Monaco et al.[93] and Zheng et al.[94]): We computed the distance to the maze at each time by taking the minimum of the Euclidean distances from the current position to the maze samples. The maze samples were provided by the linearization algorithm[95]. We then identified periods when the distance exceeded 5 cm and extended the window in both directions until the distance dropped below 1 cm. Additionally, we required that the Euclidean distance between the start and end positions of each epoch be ≤ 15 cm, ensuring the animal remained relatively stationary during the headscan. We further required that the angular velocity be ≥ 6 rad/s. Angular velocity was computed from 2D position kinematics. We differentiated vx and vy to obtain acceleration, smoothed with 0.1 standard deviation, and then estimated instantaneous angular velocity as:

$$\omega(t) = \frac{|v_x a_y - v_{xy} a_x|}{v_x^2 + v_y^2},$$

excluding samples with near-zero speed. Locomotion was defined as periods when speed was ≥ 5cm/s and not during headscans.

**Data preprocessing and synchrony event definition**

We adapt the synchrony event definition from Yang et al.[96]. First, spikes from the population were binned by 2 ms and smoothed with a Gaussian filter (7.5 ms standard deviation), and Z-scored. Second, windows where the population spike counts exceed 3 standard deviations were kept and extended in both directions to when the spike counts fall back to the mean. Third, windows shorter than 50 ms and longer than 500 ms were discarded. These define the windows of population burst events (PBEs). Next, from the detected SPW-R windows, we merge windows with inter-event intervals <200 ms. Finally, we take the union of the PBE and the merged SPW-R windows to form our final list of synchrony events. Only putative pyramidal cells are included in all the decoding analyses.

#### State-space decoding of position with switching movement dynamics

We decoded the animal's 2D position from population spike counts using a state-space decoder with switching movement dynamics, following[44,97]. Let $t \in \{1, \dots, T\}$ index time bins of width $\Delta t = 20$ ms, and let $n \in \{1, \dots, N\}$ index neurons. We denote the population spike-count vector at time $t$ by

$$y_t^{(1:N)} = (y_t^{(1)}, \dots, y_t^{(N)}) \in \mathbb{N}^N, \tag{1}$$

and the full observation sequence by $y_{1:T}^{(1:N)} = \{y_t^{(1:N)}\}_{t=1}^T$.

The latent position state consists of (i) a position variable $x_t$ and (ii) a discrete dynamics variable $I_t$ indicating whether motion is locally continuous or fragmented:

$$x_t \in \mathscr{X} \subset \mathbb{R}^2, \qquad I_t \in \{\text{continuous, fragmented}\}. \tag{2}$$

For inference, the 2D environment was discretized into a grid $\mathscr{X}$ with 3 cm spatial bins; all probability distributions over $x_t$ below are defined on this discrete grid. Only putative pyramidal cells were included in decoding.

*Rate map estimation*

For each neuron $n$, we estimated a 2D firing-rate map (place field) $\lambda^{(n)}(x)$ using a kernel density estimator (KDE) with Gaussian kernel standard deviation $\sigma = 3$ cm, using only time points when the animal speed exceeded 2.5 cm/s. During decoding we evaluate $\lambda^{(n)}(x)$ on the same 3 cm grid $\mathscr{X}$.

*Observation model*

We assumed conditionally independent Poisson spiking given position:

$$p\left(y_t^{(1:N)} \mid x_t\right) = \prod_{n=1}^{N} p\left(y_t^{(n)} \mid x_t\right), \tag{3}$$

with

$$p\left(y_t^{(n)} \mid x_t\right) = \frac{\exp\left(-\Delta t\, \lambda^{(n)}(x_t)\right) \left(\Delta t\, \lambda^{(n)}(x_t)\right)^{y_t^{(n)}}}{y_t^{(n)}!}. \tag{4}$$

Equivalently,

$$p\left(y_t^{(1:N)} \mid x_t\right) = \prod_{n=1}^{N} \frac{\exp\left(-\Delta t\, \lambda^{(n)}(x_t)\right) \left(\Delta t\, \lambda^{(n)}(x_t)\right)^{y_t^{(n)}}}{y_t^{(n)}!}. \tag{5}$$

*State dynamics with switching modes*

The discrete dynamics variable $I_t$ determines the transition prior for position. In the *continuous* mode, position follows an isotropic Gaussian random-walk prior; in the *fragmented* mode, position is uniform over the environment grid:

$$p(x_t \mid x_{t-1}, I_t = \text{continuous}) \propto \exp\left(-\frac{\|x_t - x_{t-1}\|^2}{2\sigma_{\text{rw}}^2}\right), \tag{6}$$

$$p(x_t \mid x_{t-1}, I_t = \text{fragmented}) = \frac{1}{|\mathscr{X}|}. \tag{7}$$

We set the random-walk step scale to $\sigma_{\text{rw}} = 10$ cm per 20 ms bin (i.e., $\Delta t = 20$ ms).

The discrete dynamics state $I_t$ evolved as a sticky two-state Markov chain with stay probability 0.98 and switch probability 0.02.

*Causal posterior (filtering)*

Starting from a uniform initial distribution over $(x_0, I_0)$, we computed the causal posterior recursively:

$$p(x_t, I_t \mid y_{1:t}^{(1:N)}) \propto p(y_t^{(1:N)} \mid x_t)\, p(x_t, I_t \mid y_{1:t-1}^{(1:N)}), \tag{8}$$

$$p(x_t, I_t \mid y_{1:t-1}^{(1:N)}) = \sum_{x_{t-1} \in \mathscr{X}} \sum_{I_{t-1}} p(x_t \mid x_{t-1}, I_t)\, p(I_t \mid I_{t-1})\, p(x_{t-1}, I_{t-1} \mid y_{1:t-1}^{(1:N)}). \tag{9}$$

*Acausal posterior (smoothing)*

We computed the acausal (smoothed) posterior $p(x_t, I_t \mid y_{1:T}^{(1:N)})$ via a backward recursion that combines the causal posterior with a correction term from future observations. For $t = T-1, T-2, \ldots, 1$:

$$p(x_t, I_t \mid y_{1:T}^{(1:N)}) = p(x_t, I_t \mid y_{1:t}^{(1:N)}) \sum_{x_{t+1} \in \mathscr{X}} \sum_{I_{t+1}} \frac{p(x_{t+1} \mid x_t, I_{t+1})\, p(I_{t+1} \mid I_t)}{p(x_{t+1}, I_{t+1} \mid y_{1:t}^{(1:N)})}\, p(x_{t+1}, I_{t+1} \mid y_{1:T}^{(1:N)}), \tag{10}$$

where the one-step predictive distribution is

$$p(x_{t+1}, I_{t+1} \mid y_{1:t}^{(1:N)}) = \sum_{x_t \in \mathscr{X}} \sum_{I_t} p(x_{t+1} \mid x_t, I_{t+1})\, p(I_{t+1} \mid I_t)\, p(x_t, I_t \mid y_{1:t}^{(1:N)}). \tag{11}$$

We initialize the recursion at $t = T$ with $p(x_T, I_T \mid y_{1:T}^{(1:N)})$ given by the final filtered distribution.

*Marginal posterior over position and decoded trajectory*

We obtained the marginal posterior over position by summing out the dynamics variable:

$$p(x_t \mid y_{1:T}^{(1:N)}) = \sum_{I_t \in \{\text{continuous,fragmented}\}} p(x_t, I_t \mid y_{1:T}^{(1:N)}). \tag{12}$$

When a point estimate was required, we used the MAP decoded position

$$\hat{x}_t = \arg\max_{x \in \mathscr{X}} p(x_t = x \mid y_{1:T}^{(1:N)}). \tag{13}$$

**Shuffle tests to determine whether an event is ”on-manifold”**

We used two shuffle controls to build null distributions for the state-space decoder’s marginal likelihood and to isolate what structure the evidence depends on. Neuron-ID shuffling permutes the mapping between spike trains and neuron identities while keeping the same ratemap templates, testing whether high likelihood requires the correct neuron–template correspondence. Independent circular shifting rotates each neuron’s spike train by a random offset, preserving per-neuron firing statistics but destroying across-neuron co-activity, testing whether likelihood gains reflect coordinated ensemble structure rather than rate alone. If an event’s marginal likelihood exceeds the 95-percentile in both shuffles, we call it ”on-manifold” (otherwise ”off-manifold”). The marginal likelihood is given as a side product of the causal filtering in sec. Causal posterior (filtering):

$$p(y_{1:t}) = p(y_{1:t-1}) \sum_{x_t} \sum_{I_t} p(y_t^{(1:N)} \mid x_t)\, p(x_t, I_t \mid y_{1:t-1}), \tag{14}$$

where the start of the recursion is $p(y_1) = \sum_{x_t} p(y_1^{(1:N)} \mid x_t)$.

We also computed a simpler marginal likelihood based on the Naive Bayes decoder, ignoring the temporal priors:

$$p(y_{1:T}) = \sum_{t} \sum_{x_t} p(y_t^{(1:N)} \mid x_t), \tag{15}$$

and the fraction of on-manifold events did not change much.

All the subsequent analysis on replay contents were performed on the ”on-manifold” events.

**Replay metrics**

Replay metrics were computed within each continuous segment within each event, and aggregated within an event. Continuous segments were defined as consecutive time bins where $p(I_t = \text{continuous}) \geq 0.8$. A segment was further split wherever there was a jump in Euclidean distance larger than 10cm across two consecutive time bins. **Displacement** was defined as the maximal Euclidean distance between each time bin and the start of a continuous segment, and the max was taken across the segments within one event. **Displacement per sqrt(time bin)** is the normalized version, i.e.: **Displacement**$/\sqrt{\#\text{ time bins}}$. Drawing on the theoretical framework of replay as a random walk[32], we use this metric to isolate the influence of ripple frequency from the $\sqrt{T}$ scaling predicted by duration-based displacement. **Duration** was the elapsed time of each continuous segment within one event, median across the segments. **Path length** was the cumulative Euclidean distance between consecutive time points within each continuous segment within one event, median across the segments. **Speed** was defined as the path length divided by duration within each continuous segment, median across the segments within one event. We found the displacement to be a more distinguishing feature separating replay of clear locomotion trajectories from non-trajectories (e.g. where the decoded position could move back-and-forth within a confined region, which produces a high speed without directional locomotion).

**Clustered - Solo Preference Index**

The Clustered - Solo Preference Index measures the degree to which a variable is more dominant during cSPW-Rs (positive, max +1) or during sSPW-Rs (negative, min -1). Denote the variables of interest averaged within cSPW-Rs as '$c$ and within sSPW-Rs as $s$. We computed clustered-solo preference index (**Figure 4D and G**) by $(c - s)/(c + s + 2\epsilon)$. Epsilon is a small number to stabilize the variable. In **Figure 4D**, the quantity was the posterior probability of each spatial bin during NREM sleep, and $\epsilon$ was the the 5-percentile among all spatial bins of the time-averaged posterior probability during cSPW-Rs. In **Figure 4G** the variable is the time-averaged posterior probability of latent bins (summed within each category), and $\epsilon$ is set to 0.

**Unsupervised Reactivation Analysis with Jump Latent Variable Model**

To reveal reactivation contents that may not correspond to the animal's physical position, we used the Jump Latent Variable Model (JumpLVM)[45] Unlike supervised decoders, the JumpLVM learns both the latent trajectory $x_t$ and the neural tuning curves $\{r^{(n)}(x)\}_n$ with respect to the latent coordinate/bin in an unsupervised manner.

*Generative Model*

The model assumes that neural activities $Y$ are generated by a 1D latent variable $x_t$ (discretized into $\{1, \ldots, L\}$, $L$ set to 100) via smooth tuning curves. To ensure smoothness, we model the tuning curves as Gaussian Processes (GP) using a basis function expansion. The generative process is defined as:

$$W_{bn} \sim \text{Normal}(0, \sigma_w^2) \tag{16}$$

$$r_{tn} = g\left(\sum_b \phi_b(x_t) W_{bn}\right) \tag{17}$$

$$y_{tn} \sim \text{Poisson}(r_{tn}) \tag{18}$$

where $g(\cdot)$ is the Softplus non-linearity. The basis functions $\phi_b$ are derived from the spectral decomposition of the RBF kernel matrix $\mathbf{K}$, where $\mathbf{K}_{ij} = \exp\left(-\frac{(i-j)^2}{2l^2}\right)$. Specifically, $\phi_b = \sqrt{\lambda_b}\mathbf{v}_b$, where $\lambda_b$ and $\mathbf{v}_b$ are the eigenvalues and eigenvectors of $\mathbf{K}$. We set $\sigma_w^2 = 1, l = 10$.

*Latent Dynamics*

The latent follows the same dynamics as in the state-space decoder in sec. State-space decoding of position with switching movement dynamics. We used a movement variance of 1 and probability of staying in the same dynamics 0.99.

*Inference via EM Algorithm*

We maximize the log marginal posterior,

$$L(W, \theta) = \log \int p(Y, X|W, \theta)p(X|\theta)dX + \log p(W|\theta), \tag{19}$$

using the Expectation-Maximization (EM) algorithm[99]. $\theta$ denotes the hyperparameters, including the prior variance $\sigma_w^2$ of the weights, lengthscale $l$ of the GP kernel, movement variance of the latent random walk, and the transition probability of the discrete dynamics types (see sec.State-space decoding of position with switching movement dynamics).

**E-step (State-Space Decoding):** We compute the acausal posterior $q_t^* = p(x_t \mid Y, W^*, \theta)$ using forward-backward smoothing recursions and maginalizing over the dynamics $I_t$, same as in sec. Acausal posterior (smoothing).

**M-step (Tuning Curve Estimation):** We update the weights $W$ by maximizing the expected complete-data log-posterior (LCD):

$$LCD(W|\theta) = \mathbb{E}_{q^*}[\log p(Y, X|W, \theta)] + \log p(W|\theta). \tag{20}$$

This step is conceptually similar to tuning curve estimation with a generalized linear model[100], although the regressor now has uncertainty that needs to be averaged over. After some unpacking and derivations (see[45]) , the objective is computed efficiently using summary statistics:

$$LCD(W \mid \theta) \sim \sum_l \sum_n (s_{ln} \log r_{ln} - c_l r_{ln}) + \log p(W \mid \theta) \tag{21}$$

where $c_l = \sum_t p(x_t = l \mid Y, W^*, \theta)$ is the effective occupancy and $s_{ln} = \sum_t p(x_t = l \mid Y, W^*, \theta) y_{tn}$ is the effective spike count. This optimization is performed using the ADAM optimizer.

We initialized the posterior $q^* = p(X|Y, W, \theta)$ as uniform (i.e. $= 1/L$) plus a small random perturbation. The perturbation for each latent bin and time bin was sampled independently from a uniform distribution between $[0, 0.1]$. We then re-normalized it to sum to 1 in the latent dimension. We then iterated the two EM steps for 20 iterations (when convergence was achieved).

*Decode synchrony events*

We trained the model on spike counts binned at 100ms during the task epoch. After training, we decoded the posterior probability of the latent during the synchrony events during NREM sleep using a naive Bayes decoder to avoid potential bias from the latent transition priors:

$$p(x_t = i|y_t^{(1:N)}, W) = \frac{p(y_t^{(1:N)}|x_t = i, W)}{\sum_j p(y_t^{(1:N)}|x_t = j, W)}, \tag{22}$$

which also gives the marginal likelihood of each event:

$$p(Y_{event}|W) = \sum_{t \in \text{event}} \sum_j p(y_t^{(1:N)}|x_t = j, W). \tag{23}$$

The likelihood $p(y_t^{(1:N)}|x_t = i, W)$ depends on the tuning curves learned from the task epoch. To decode the sleep events we multiplied the tuning curves by a gain factor greater than 1. The gain factor was chosen for each session to maximize the fraction of significant events (same shuffle tests as in sec. Shuffle tests to determine whether an event is ”on-manifold”) among half of the events (randomly selected). (We used a gain of 1 in the supervised state-space decoder case, because sweeping the gain did not produce much change in the fraction of significant events. Furthermore, for each gain value we tested, the conclusion that the latent variable model detected more on-manifold events holds. **Figure S15**)

Similar to analyzing with the supervised decoder as stated in sec. Shuffle tests to determine whether an event is ”on-manifold”, only significant events (”significant” relative to the latent tuning curves) were used for downstream latent analysis.

#### Behavior type scoring and classification of latent bins

**Latent behavioral scoring and classification.** To score and classify the latent bins based on behavior types, we computed the mean posterior mass of latent $k$ within each behavior type $b \in$ {immobility, locomotion, headscan}:

$$\bar{p}_{i,b} = \frac{1}{T_b} \sum_{t \in b} p(x_t = i \mid Y),$$

where $T_b$ is the number of time bins assigned to behavior type $b$. We then normalized across behavior types for each latent to obtain a *behavior score*:

$$f_{k,b} = \frac{\bar{p}_{k,b}}{\sum_{b'} \bar{p}_{k,b'}}.$$

For instance, if a latent occurs in both locomotion and headscan, but out of a total of 10 time bins from headscan it occured 9 times, and out of a total of 1000 time bins from locomotion it also occured 9 times, it would have a higher score for headscan. A latent $k$ was labeled as behavior type $b$ if $f_{k,b} \geq 0.5$; latents with $f_{k,b} \leq 0.5$ for all $b$ were unclassified.

#### Statistical analysis

Statistical analysis included 2 x 2 repeated-measures ANOVA with planned contrasts and Wilcoxon signed-rank and rank-sum tests. The ANOVA was performed using statsmodels.stats.anova.AnovaRM. P-values from the contrasts were adjusted using the Holm method via statsmodels.stats.multitest.multipletests. The Wilcoxon signed-rank and rank-sum tests were performed via scipy.stats.

### Supplementary Figures

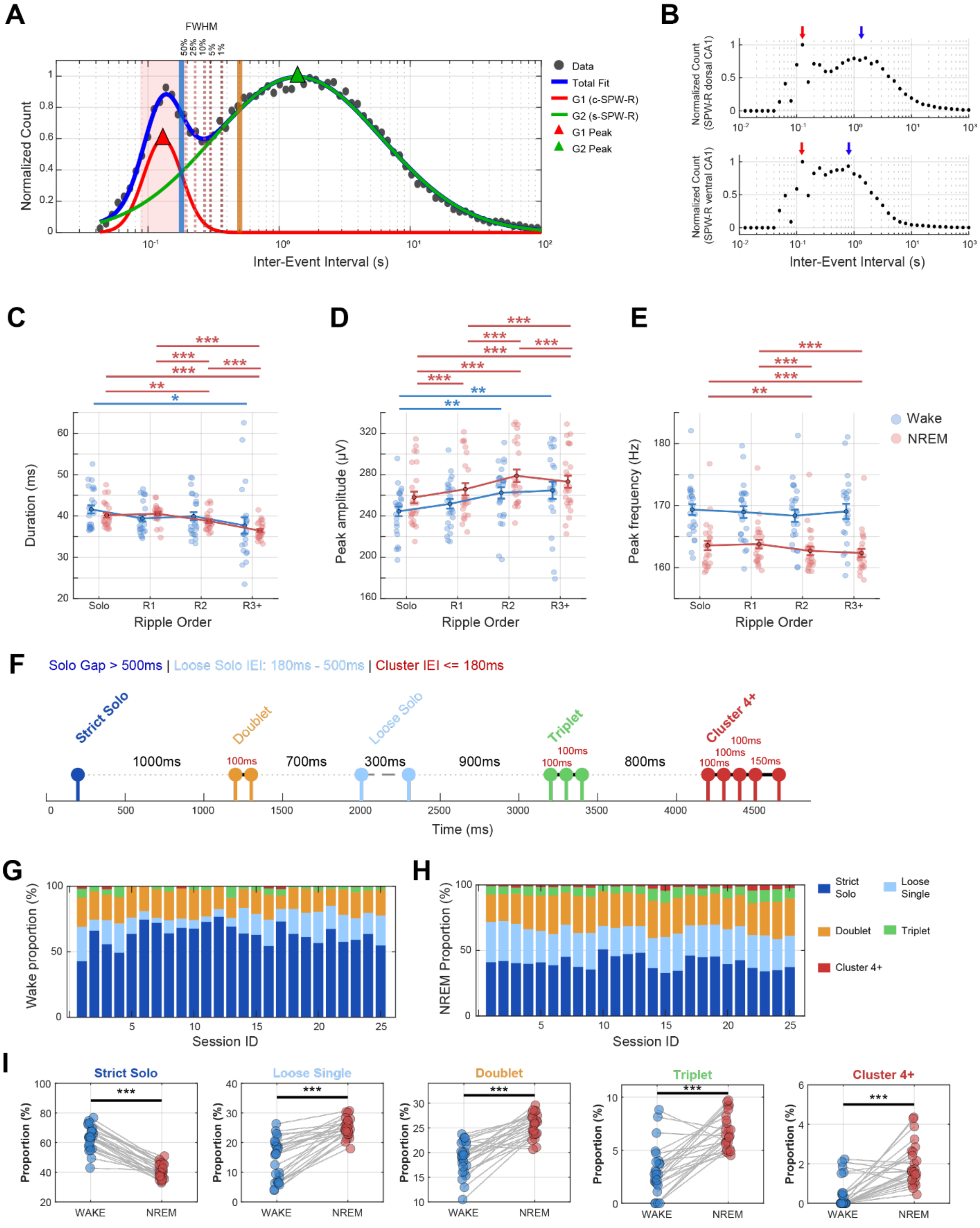

**Figure S1. Prevalence and state-dependent modulation of ripple clusters.** Related to Fig. 1. **(A)** Inter-event interval (IEI) distribution of ripples across all sessions (n = 86354 ripples from 25 sessions, 7 mice) fitted with a double Gaussian model. Data points (black circles) represent binned IEI counts on a logarithmic scale. The total fit (blue solid line) comprises two distinct Gaussian components: Gaussian 1 (red line, "cluster" population, centered at 130 ms, FWHM: 87-193 ms) and Gaussian 2 (green line, "solo" population, centered at 1383 ms, FWHM: 242-7915 ms). Vertical dotted lines mark decay thresholds where Gaussian 1 falls to specific percentages of its peak amplitude (50%, 25%, 10%, 5%, 1%). Red shaded region indicates the full-width at half-maximum (FWHM) of the cluster distribution. Solid vertical lines denote empirically validated thresholds: 180 ms for electrophysiology (cyan) and 500 ms for widefield $Ca^{2+}$ imaging (orange). **(B)** Longitudinal IEI distributions from three rats with dorsal-intermediate or dorsal-ventral recordings. Dorsal distribution peaks: 126 ms and 1585 ms. Combined intermediate/ventral peaks: 126 ms and 794 ms. **(C-E)** Ripple properties across Solo (>500 ms isolation) and cluster positions (R1, R2, R3+; classified using 180 ms threshold). Statistics: Linear Mixed Effects models with post-hoc FDR correction. **(C)** Duration: WAKE showed minimal variation; NREM showed progressive shortening within clusters (R1 → R2 → R3+, all $p < 0.001$). **(D)** Peak amplitude: WAKE solo ripples smaller than R2/R3+ ($p < 0.002$). NREM showed systematic increases from Solo → R1 → R2, then decreased at R3+. **(E)** Peak frequency: stable across categories in WAKE; NREM showed progressive decrease (Solo/R1 > R2/R3+, $p < 0.001$). **(F)** Classification schematic: clusters defined by ≤180 ms IEI (Doublets, Triplets, ≥4). Solo events: Strict (>500 ms isolation) or Loose (180-500 ms, "grey zone"). **(G-H)** Stacked bar charts showing the rate of ripple events (Solo, Doublet, Triplet, ≥4) per session during WAKE **(G)** and NREM sleep **(H)**. Each vertical bar represents one session. **(I)** Paired t-tests on proportions (n = 25 sessions) revealed: Strict Solo (WAKE > NREM, $p < 0.001$), Doublet, Triplet and Cluster 4+ (NREM > WAKE, $p < 0.01$). The cSPW-R/sSPW-R ratio increased ~2-fold from WAKE (1.2-1.5) to NREM (2.5-3.0), suggesting NREM-specific mechanisms promote sustained cluster generation.

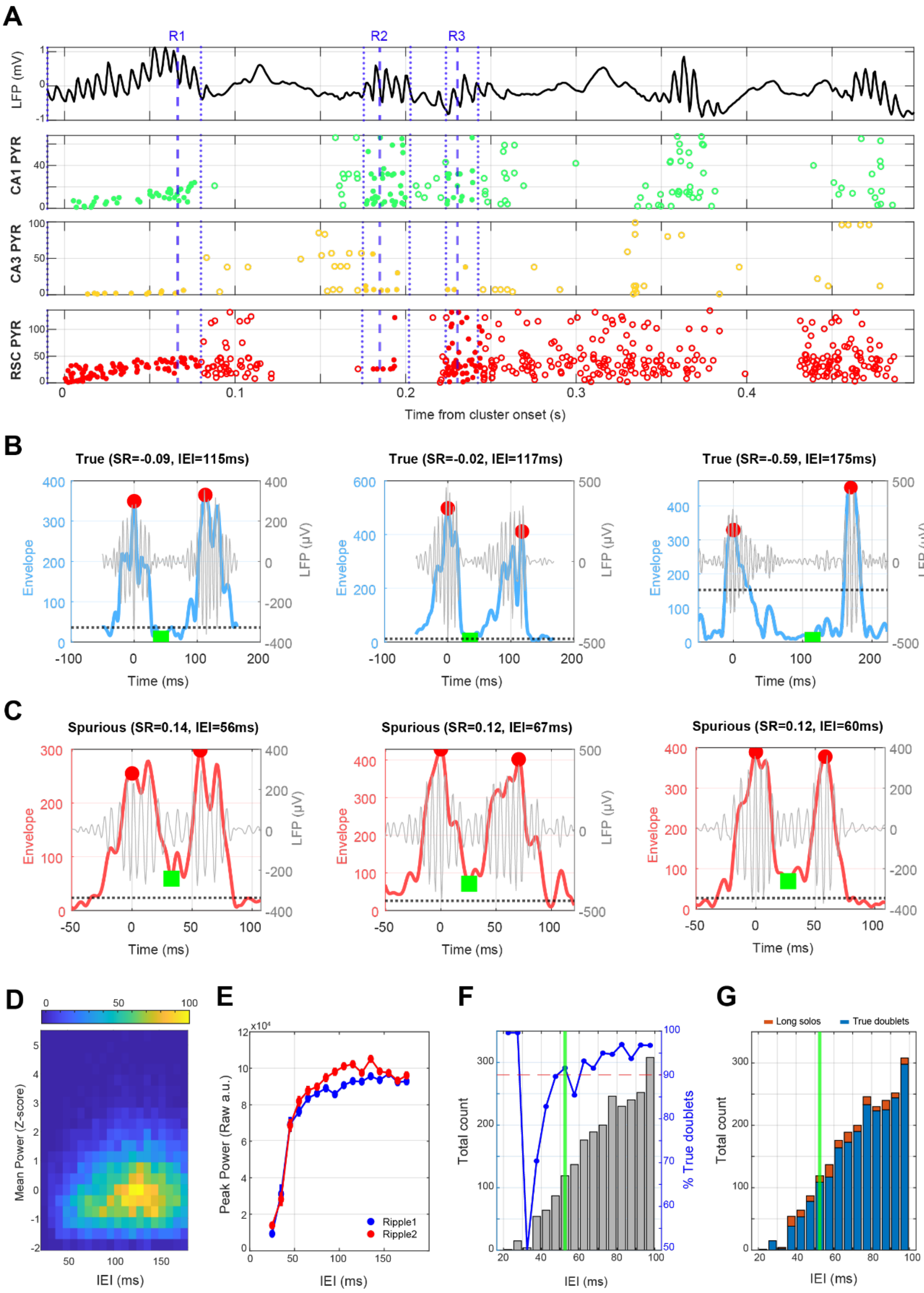


**Figure S2. Differentiation of Ripple Clusters and Spurious Single "Split" Events.** Related to Fig. 1. Representative examples illustrating the morphological criteria used to filter artifacts from the dataset. (**A**) Example trace. Wideband LFP signal of ripples (top) showing clustered ripple events labeled R1, R2, R3 and two isolated SPW-Rs. Bottom three panels: Firing of CA1 pyramidal cells (PYR, green), CA3 pyramidal cells (yellow), and retrosplenial cortex (RSC, red) neurons. Note the continuous firing of CA1 pyramidal cells between R2 and R3, suggesting these may represent a single prolonged event (long solo ripple, sSPW-R) instead of two distinct events. **(B)** Three examples of valid doublet ripples. The smoothed Hilbert envelope (thick blue line) exhibits a pronounced dip between the two ripple peaks (red circles), returning near the baseline (black dotted line) before the second event is initiated. This "deep saddle" (green) separates two distinct oscillation events. **(C)** Three examples of spurious doublets (false detections), where the envelope (blue) remains well above baseline between peaks (green). Gray traces = band-pass filtered LFP (120–250 Hz). Red circles = detected local maxima. Green squares = saddle point (inter-peak minimum). Black dotted line = calculated baseline amplitude. Classification was performed using a "Saddle Ratio (SR)" threshold: pairs were rejected if the saddle height exceeded 10% of the peak amplitude relative to baseline. Note different time scales in B and C. **(D)** Two-dimensional density histogram (Z-scored power vs. inter-event intervals. The majority of detected doublet ripples cluster between 100-150ms inter-event intervals with near-average power. **(E)** Comparison of absolute peak power (raw arbitrary units) between the first (blue) and second (red) ripple in doublets across inter-event interval bins. Ripples show diminished power at very short intervals (<50ms). In the physiological doublet range (>50ms), the second ripple consistently exhibits higher peak power than the first. **(F)** Quantification of doublet quality as a function of inter-event intervals. Gray bars (left axis) show the total count of candidate pairs in 5-ms bins. Blue line (right axis) indicates percentage classified as true doublets based on combined power and envelope criteria. Red dashed line marks 90% threshold. Acceptance rate drops sharply below IEI=50ms. Green vertical line indicates the recommended minimum IEI threshold (53 ms), where dataset quality consistently exceeds 90%. **(G)** Classification Distribution across IEI range. Stacked bars show absolute counts of events classified as true doublets (blue) versus long solo ripples (orange). Note that the artificially "split" ripples (orange bars) are relatively disproportionate in the <50ms range. Imposing the 52.5ms cutoff (green line) effectively removes the majority of spurious double ripples. Analysis involved 2322 ripple doublets with IEIs between 20-100ms across 14 recording sessions (N=4 mice). Criteria to distinguish true doublets from 'long solos' included (1) deep power troughs (trough/peak ratio <0.1), (2) return to near-baseline power (trough/baseline ratio <1.5), and (3) absence of secondary peaks in the gap region. Using a saddle ratio threshold of 0.10, we classified 7396 of 7550 pairs (98.0%) as true doublets and only 154 (2.0%) were split artifacts across all sessions.

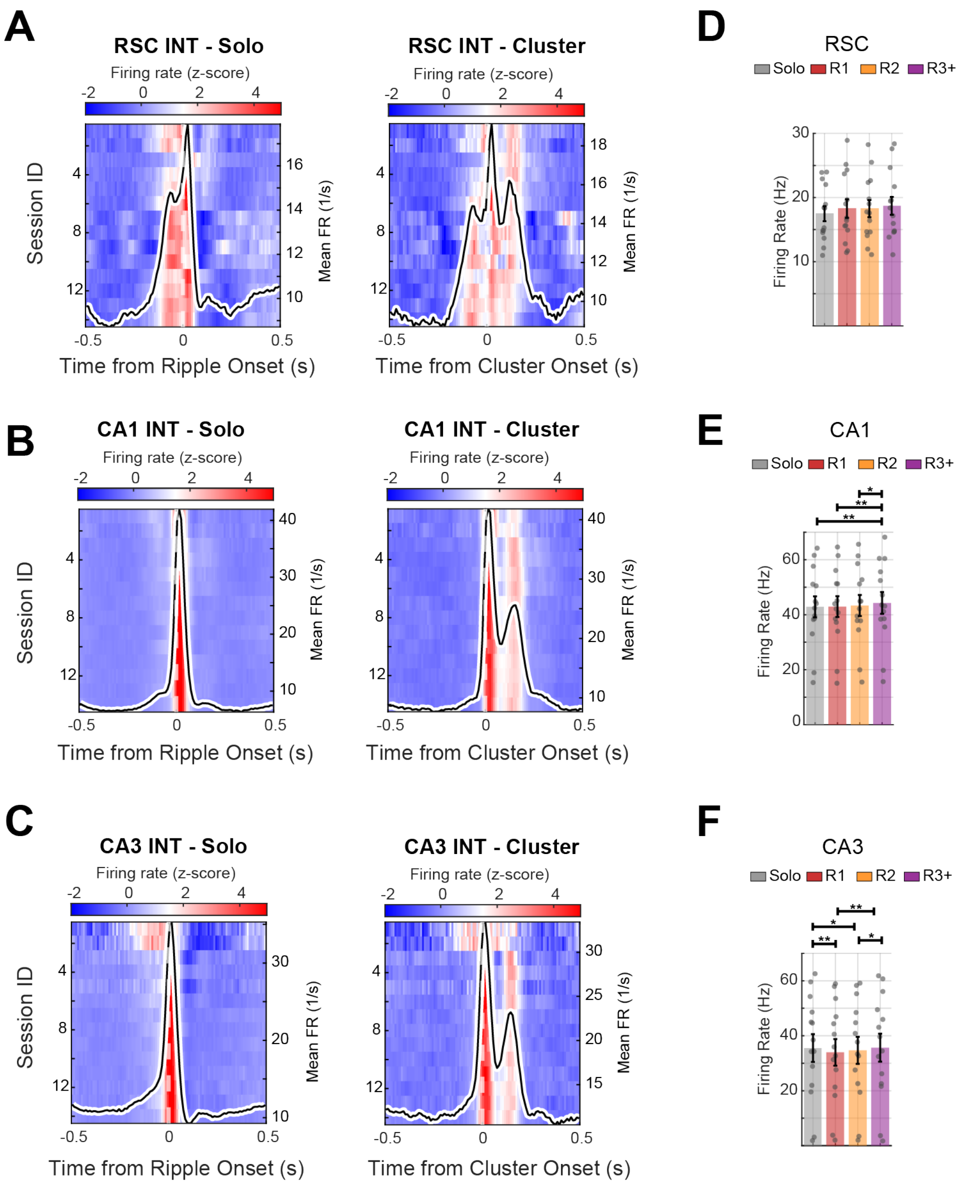


**Figure S3. Spiking rate of putative interneurons across in CA3-CA1-RSC.** Related to Fig. 1. **(A-C)** Population-average firing rates (z-scored) aligned to Solo ripple onset (left) and Cluster onset (right) for RSC **(A)**, CA1 **(B)**, and CA3 **(C)**. Heatmaps show session-averaged responses (n=14 sessions), and black traces indicate the mean firing rate across all sessions. Within each region, putative interneurons (INT) are shown. **(D-F)** Bar plots summarize average firing rates during solo ripples (blue) and the first three ripples of each cluster (R1, R2, R3; coral). Individual session averages are overlaid as grey dots. Asterisks above bars indicate statistically significant differences based on FDR-corrected linear mixed-effects (LME) models ($p < 0.05$: *, $p < 0.01$: **, for details see **Supplementary Table 3**).

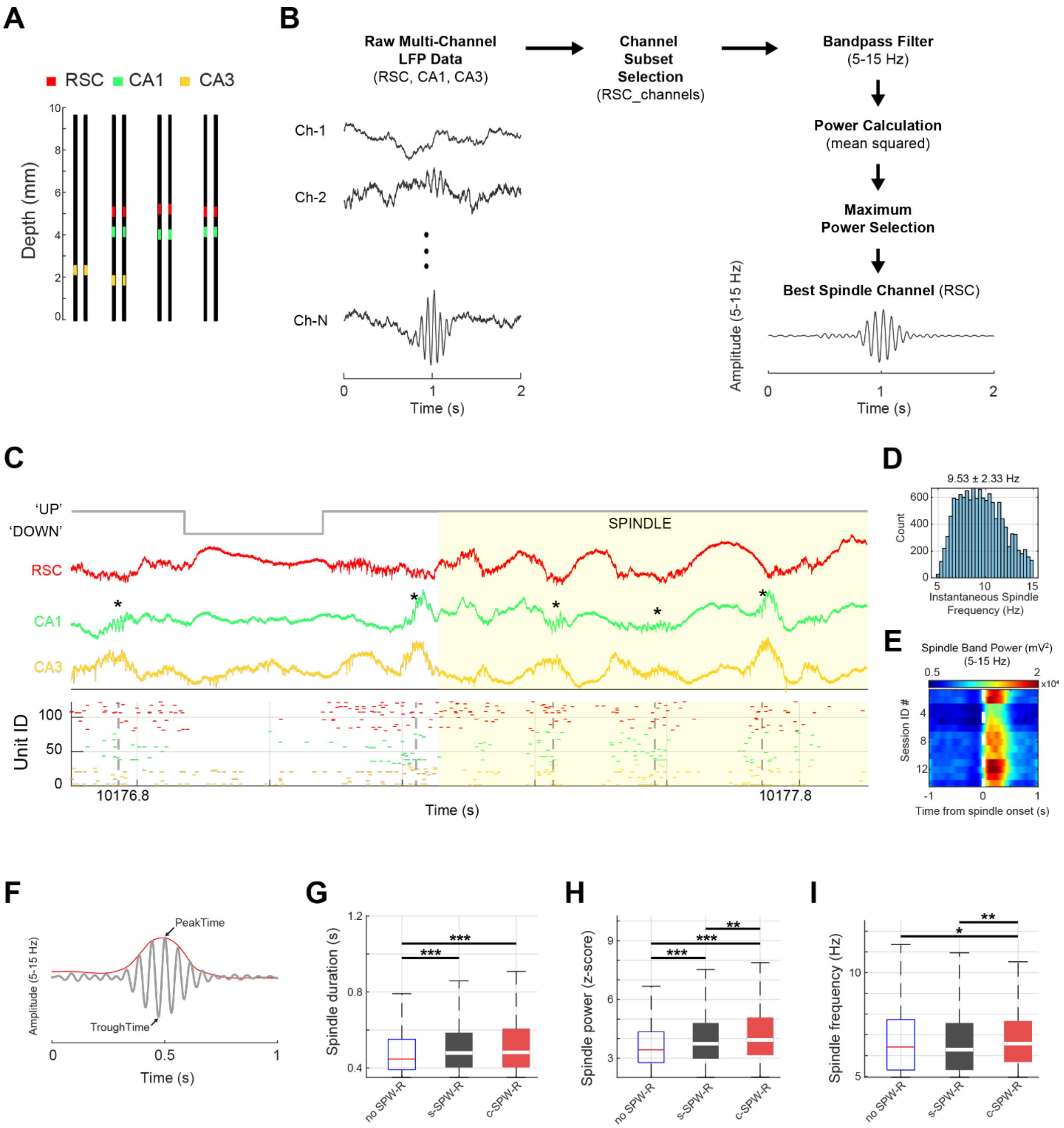


**Figure S4. Spindle properties. (A)** Schematic layout of the NP2.0 probe. Black rectangles show the recording channels of NP2.0. Black rectangles indicate recording channels; colored squares mark channels selected for simultaneous recordings from retrosplenial cortex (RSC, red), CA1 (green), and CA3 (yellow). **(B)** Spindel detection pipeline. **(C)** Wideband LFP traces from RSC (red), CA1 (green), and CA3 (yellow). Cortical UP-DOWN transitions (solid grey line) and spindle oscillations (yellow highlight, 5-15 Hz) were identified from RSC. CA1 SPW-Rs (grey dotted lines) were identified in the CA1 pyramidal layer. Bottom: raster plot shows multi-unit spiking across regions. Each row represents spikes from a putative single neuron. **(D)** Distribution of instantaneous spindle frequencies (n=14 sessions). The population mean frequency was 9.53 ± 2.33 Hz. **(E)** Heatmap of spindle-band power (5-15 Hz) aligned to spindle onset across all 14 sessions (n=4 mice), demonstrating consistent spindle detection and high signal-to-noise ratio across animals. **(F)** Schematic of spindle detection showing the filtered spindle-band oscillation (5-15 Hz, gray) and instantaneous amplitude envelope (red). Spindle trough (TroughTime) and peak amplitude time (PeakTime) are marked. **(GE)** Spindle duration as a function of ripple presence. Spindles containing solo (s-SPW-R, n=882) or clustered ripples (c-SPW-R; n=579) were significantly longer than spindles without ripples (no SPW-R, n=1882, Kruskal-Wallis with post-hoc Wilcoxon tests). **(H)** Spindle peak power (z-scored amplitude) as a function of ripple presence. Spindles with c-SPW-Rs showed the highest amplitude (no SPW-R: 3.73±1.36, s-SPW-Rs: 4.03±1.44; c-SPW-Rs: 4.31±1.63), with all pairwise comparisons being significant (Kruskal-Wallis p<0.001). **(I)** Spindle frequency shows subtle but significant differences across groups (Kruskal-Wallis p=0.003). c-SPW-R-associated spindles show higher frequency (6.87±1.56 Hz) compared to no SPW-Rs (6.71±1.59 Hz) and s-SPW-Rs (6.66±1.63 Hz) spindles.

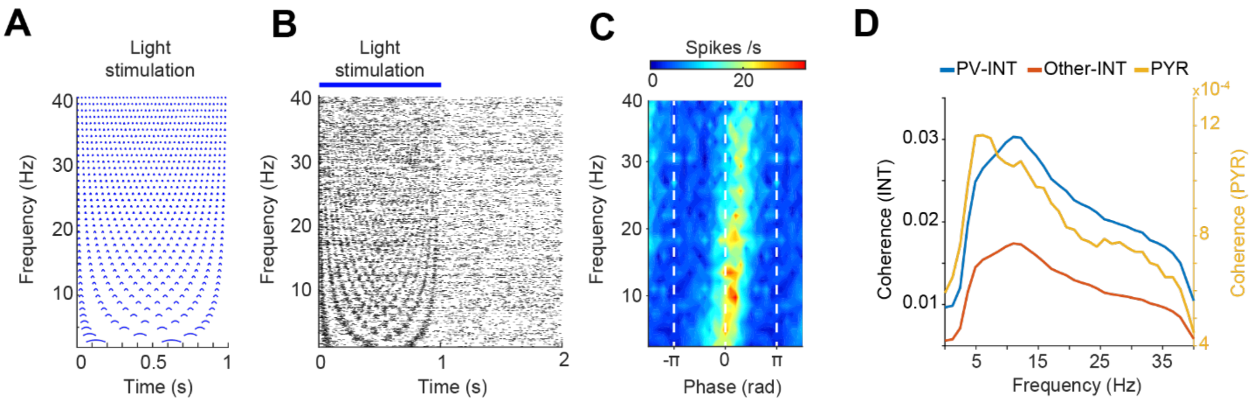


**Figure S5. Resonant properties of hippocampal parvalbumin interneurons in the theta-spindle band**. Related to Fig. 1. **(A)** Neuronal activity was recorded from the dorsal CA1 of the hippocampus with a Neuropixels 2.0 optrode in a transgenic mouse expressing ChR2 selectively in parvalbumin (PV) expressing inhibitory cells. Pure sinusoidal optical stimuli were used from 1 to 40 Hz. **(B)** Left: raster plot of an example opto-tagged PV neuron. Different frequency sinusoidal light stimulation (1 s) was followed by 1 s of no stimulation (1200 trials, 30 trials/frequency). Right: mean waveform and autocorrelation histogram of the PV neuron. Bottom: spiking gain during light stimulation. Gain plotted as a frequency-phase map (bin size: 2 Hz, π/10 rad). Spikes are evoked around the peaks of the sinusoidal signal (0 radians) at all frequencies, with the highest gain (33 spikes/s at 8 Hz stimulation). **(C)** Quantification of spiking resonance (mean) of PV positive (n = 21, blue), other putative interneurons (n = 49, red), and putative pyramidal cells (Pyr, n = 145; yellow, n = 2 mice, 3 sessions) [98]. Note the strongest resonance response between 4-15 Hz.

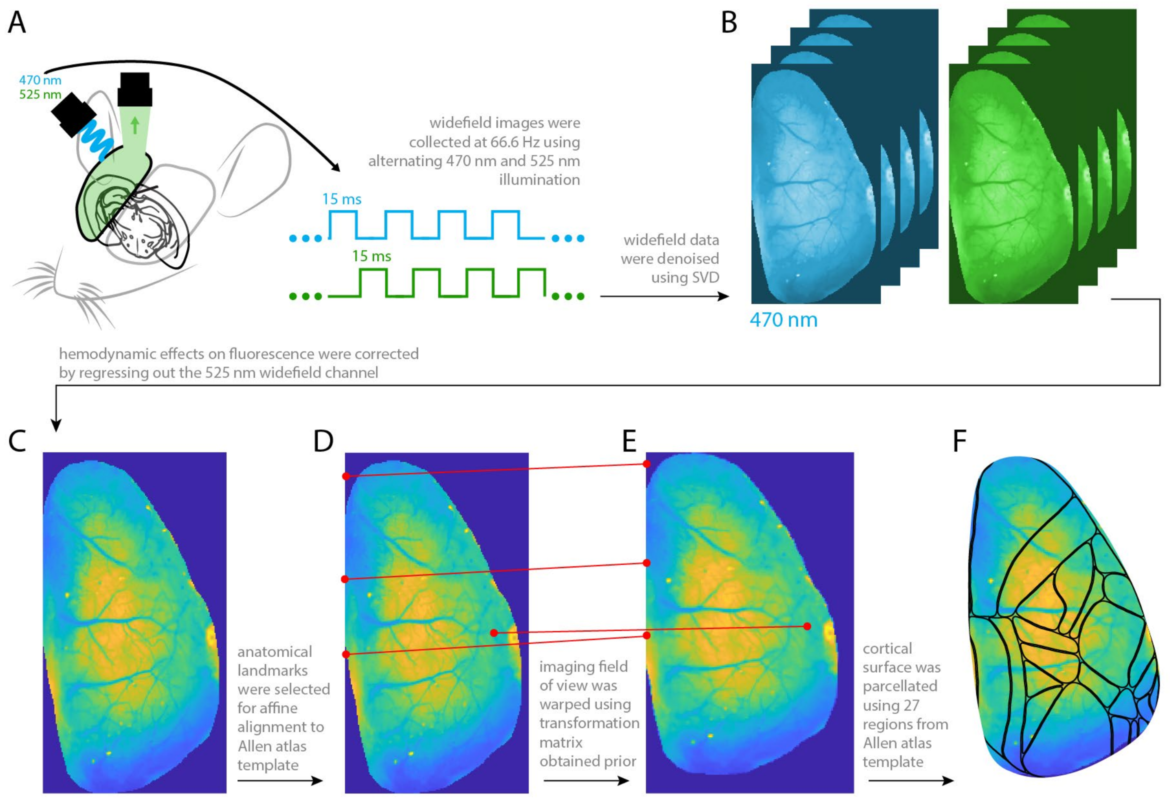


**Figure S6. Processing pipeline for widefield imaging.** Related to Fig. 2 and 3. Steps outlined in figure.

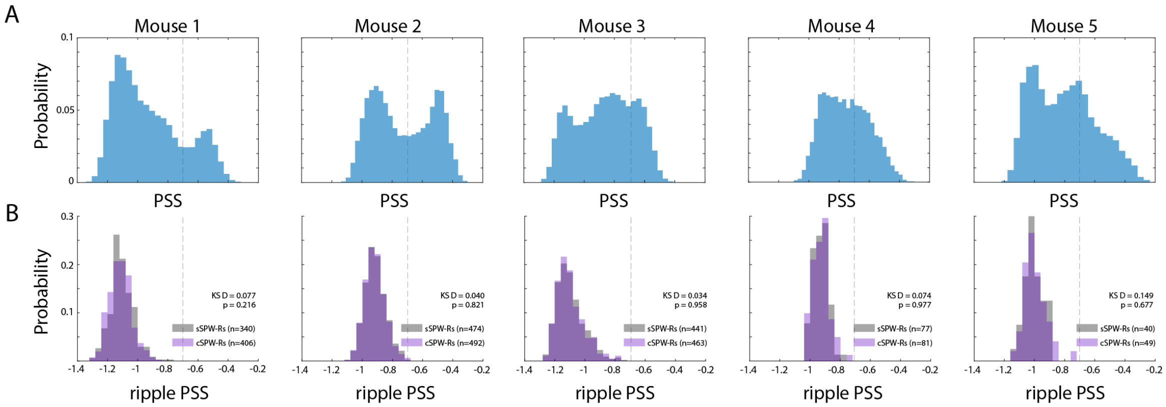


**Figure S7.** Related to Fig. 2. **(A)** Probability distribution of the slope of the power spectrum (PSS) for individual mice. **(B)** Probability distribution of SPW-R-triggered PSS values for individual mice. Following subsampling, **s**SPW-R and **c**SPW-R PSS distributions did not differ significantly (D<0.15, p >0.21 across animals). Vertical dotted line at PSS = 0.7 represents the threshold for deep NREM and ripple inclusion.

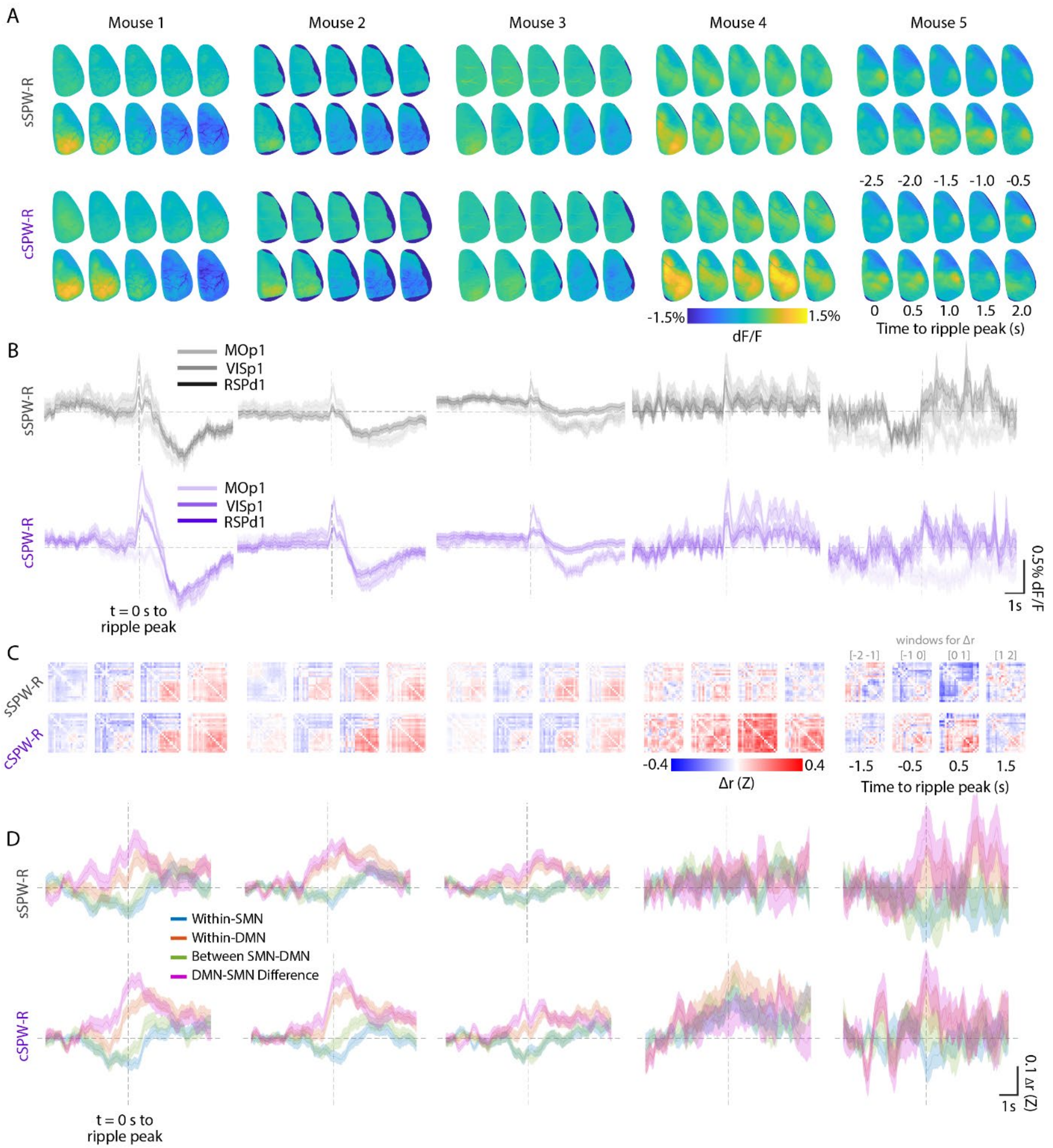


**Figure S8. Extended time course of solo and clustered ripple effects on global cortical activity.** Related to Fig. 2. All data are shown for individual mice. **(A)** Sequence of frames depicting WF activity relative to **s**SPW-R or **c**SPW-R peak at t = 0, averaged across ripples. **(B)** s/cSPW-R-triggered median fluorescence of 3 representative subregions: primary motor cortex (*MOp1*), primary visual cortex (*VISp1*), and dorsal retrosplenial cortex (*RSPd1*). **(C)** Sequence of matrices depicting s/cSPW-R-triggered change in pairwise regional correlations, averaged across ripples. SPW-R-triggered changes in pairwise regional correlations (Z) were computed using a 1 second sliding window, followed by a baseline subtraction using the -5 to -2 second range prior to ripple peak. **(D)** For default mode (DMN) and somatomotor networks (SMN), traces show average SPW-R-triggered change in within- and between-network correlations (Z). Shaded areas (**B,D**) represent SEM.

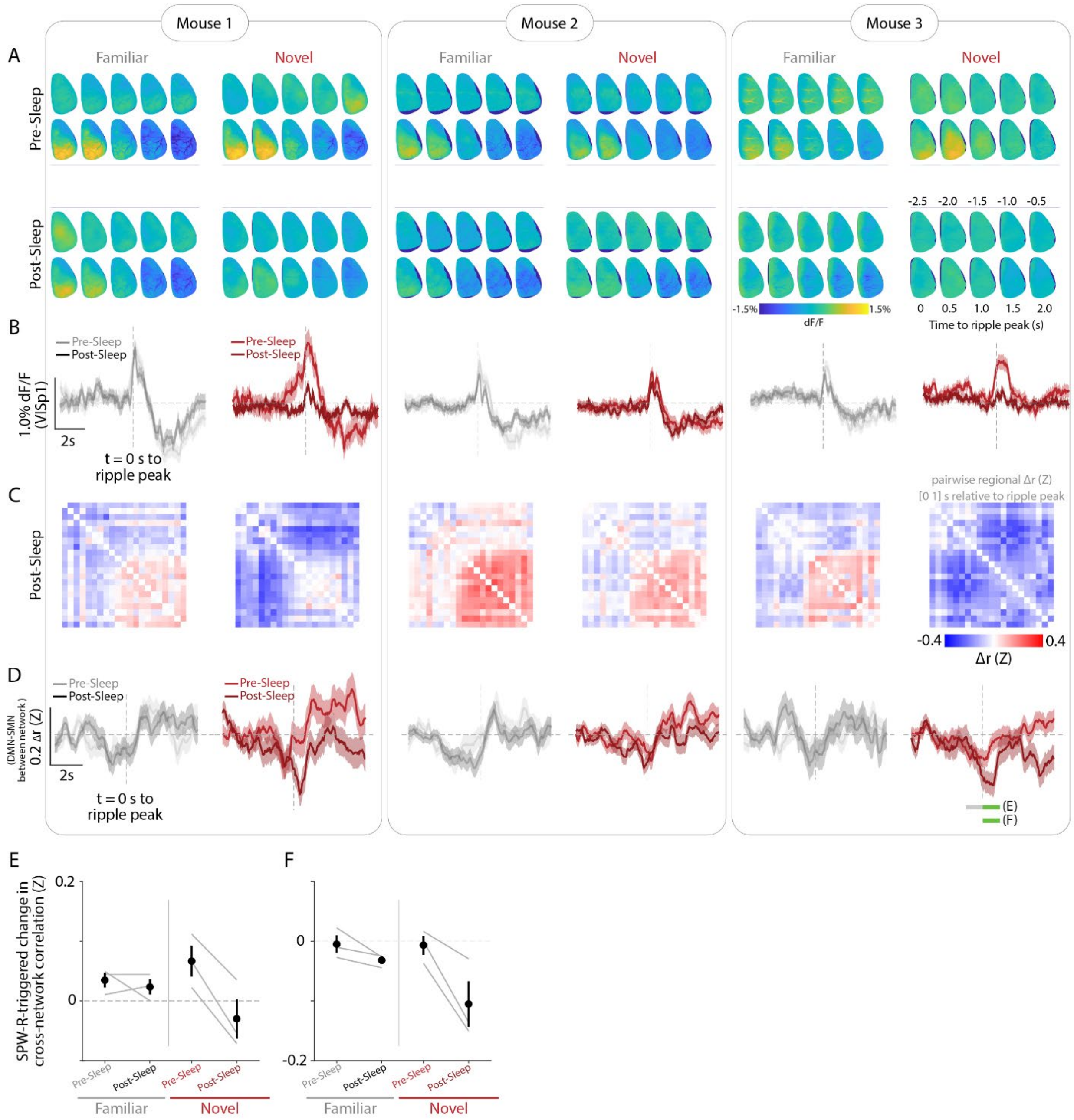


**Figure S9. Extended time course of clustered ripple effects on global cortical activity before and after familiar or novel experiences.** Related to Fig. 3. All data are shown for pre- and post-sleep conditions across behavioral contexts (familiar/novel) for individual mice. **(A)** Sequence of frames depicting WF activity relative to **c**SPW-R peak at t = 0, averaged across ripples. **(B)** **c**SPW-R-triggered median fluorescence of default motor network representative subregion (*VISp1*). **(C)** Matrix depicting **c**SPW-R-triggered change in pairwise regional correlations (Z) in the 1-s window after ripple peak. SPW-R-triggered changes in pairwise regional correlations (Z) were computed using a 1-s sliding window, followed by a baseline subtraction using the [-5 to-2] second range prior to ripple peak. **(D)** **c**SPW-R-triggered change in between-network correlations (Z) for default mode (DMN) and somatomotor networks (SMN). Green and grey bars (*rightmost panel*) represent times used for summary statistics in (E) and (F). **(E-F)** Summary of ripple-triggered DMN-SMN decorrelation was computed in **(E)** using the difference in mean traces between the timespans indicated by the green and grey bars in (D), or in **(F)** by using the mean traces obtained in the timespan indicated by the green bar alone. Post-sleep cSPW-Rs are associated with reduced between-network correlation ($F_{(1,2)} > 54.322$. $p < 0.05$) and this effect is pronounced in the novel condition ($t_{pre\ vs\ post\ (2)} = 7.61$, $p < 0.05$ for (E) and Fig. 1G; $t_{pre\ vs\ post\ (2)} = 3.63$, $p =0.07$ for (F)).

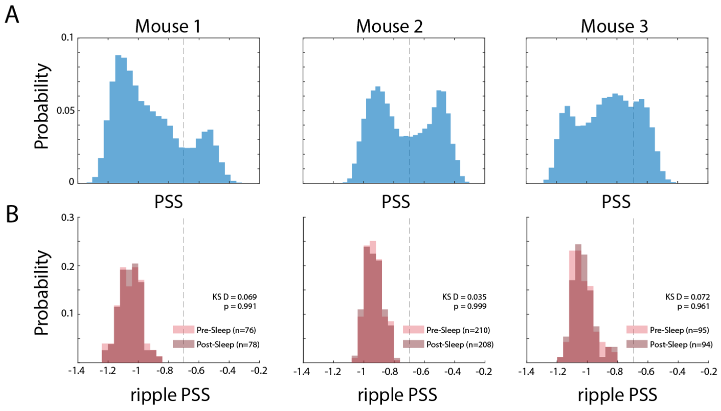


**Figure S10. The effects of clustered ripples on global cortical activity before and after novel experience were compared across similar brain states.** Related to Fig. 3. **(A)** Same as Suppl. Fig. 7A. Probability distribution of the slope of the power spectrum (PSS) across multiple sessions for individual mice. **(B)** Probability distribution of **c**SPW-R-triggered PSS values across individual mice. Following subsampling, **c**SPW-R PSS distributions before and after novel experience did not differ significantly ($D < 0.07$, $p > 0.96$ across animals). Vertical dotted line at PSS = 0.7 represents threshold for deep NREM and ripple inclusion.

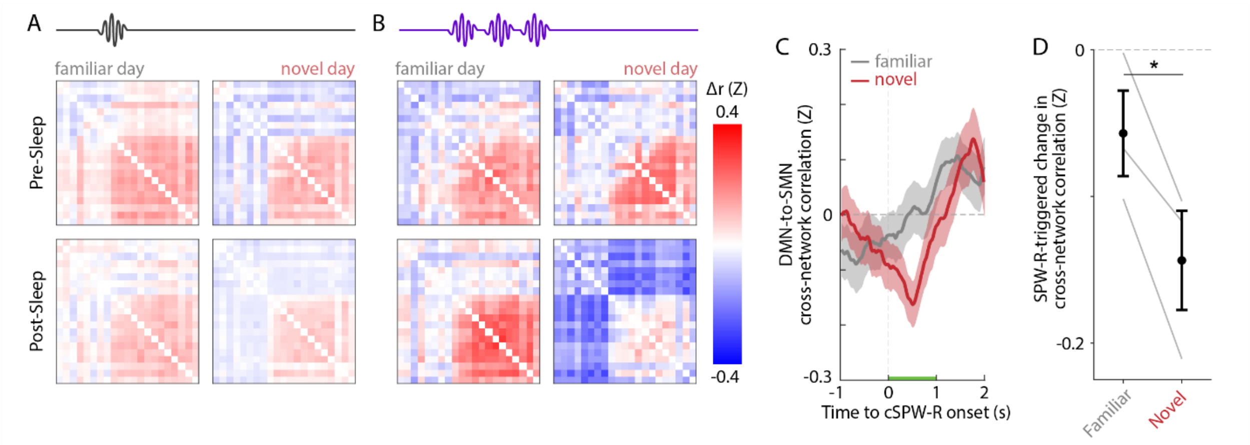


**Figure S11. The segregation effect of cSPW-Rs is consistent across cluster definitions.** Related to Fig. 3 and S9. Data are shown for **c**SPW-Rs containing 3 or more ripples separated by less than 500 ms. **(A-B)** SPW-R-triggered change in pairwise regional correlations (Z) across behavioral conditions for a representative animal (*mouse 2* in Fig. S5). Matrices represent Δr (Z) computed in the one-second window following ripple peak. **(C-D)** **c**SPW-R-triggered change in DMN-SMN between-network correlations following familiar or novel experience for the same animal and (D) across animals ($t_{(2)} = -4.67$*). Change in cross-network correlation was computed as the baseline-subtracted aggregate between-network correlation (Z-transformed) obtained in the 1-s window denoted by the green bar in (C). Shaded areas (C) and error bars represent SEM. *$p < 0.05$.

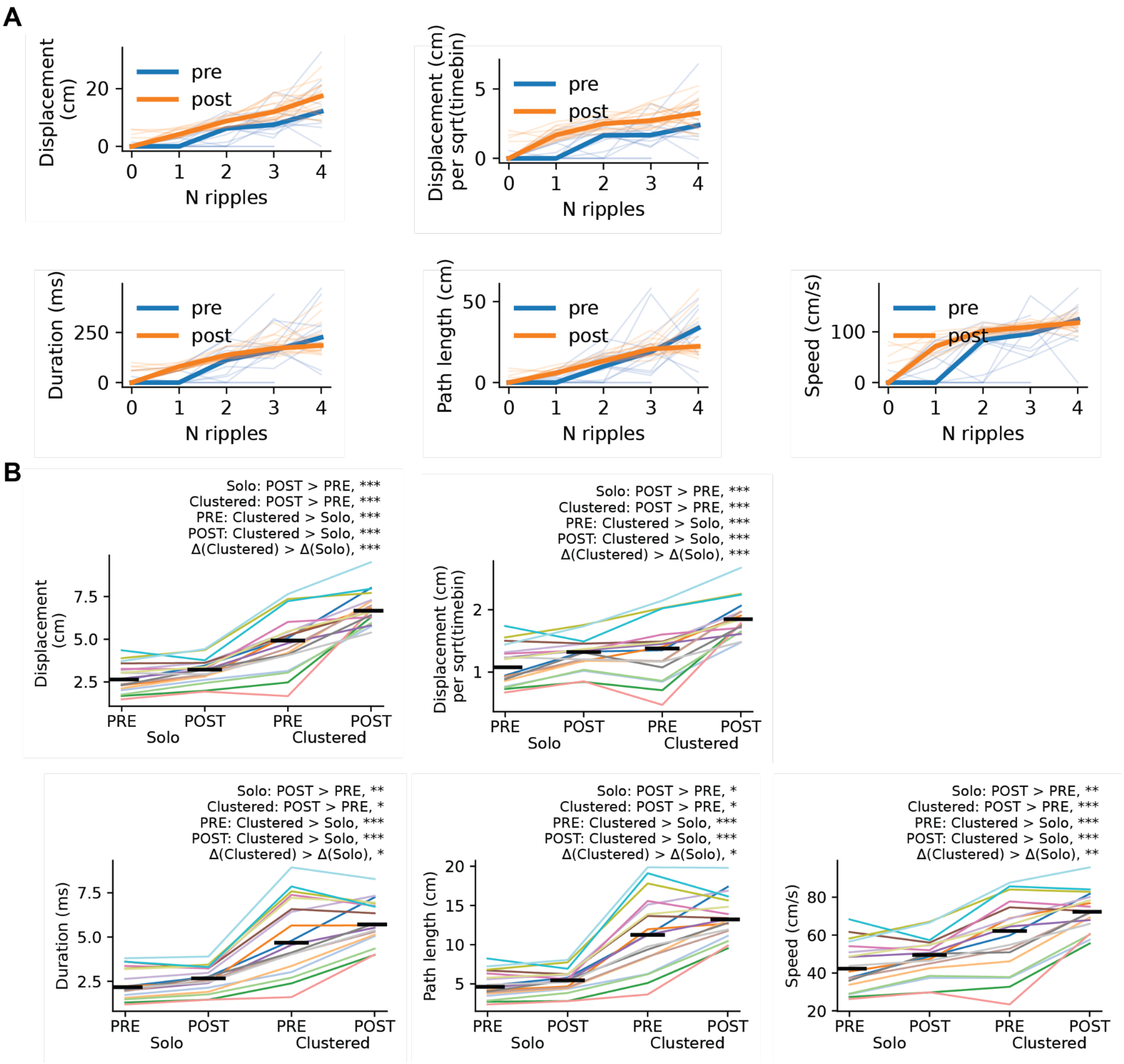


**Figure S12. Additional replay metrics and c/s SPW-R comparison.** (**A**) Metrics of replayed trajectories as a function of the number of SPW-Rs contained in a synchrony event. Each metric (Y label) is separated into PRE-task sleep (blue) and POST-task sleep (orange). Each dim line is a median within all synchrony events during PRE-task (blue) / POST-task (orange) for one session. Thicker lines are median over sessions. Note that the metrics for one event were computed separately within each continuous segment, and the median across continuous segments was computed within one event. (**B**) The values in (**A**), grouped into events that contained either **s**SPW-Rs (solo) or **c**SPW-Rs (clustered), and PRE/POST-task sleep. Each colored line is median within one session. Black bars are medians across sessions. Significance was determined via a 2 x 2 repeated measure ANOVA with planned contrasts.

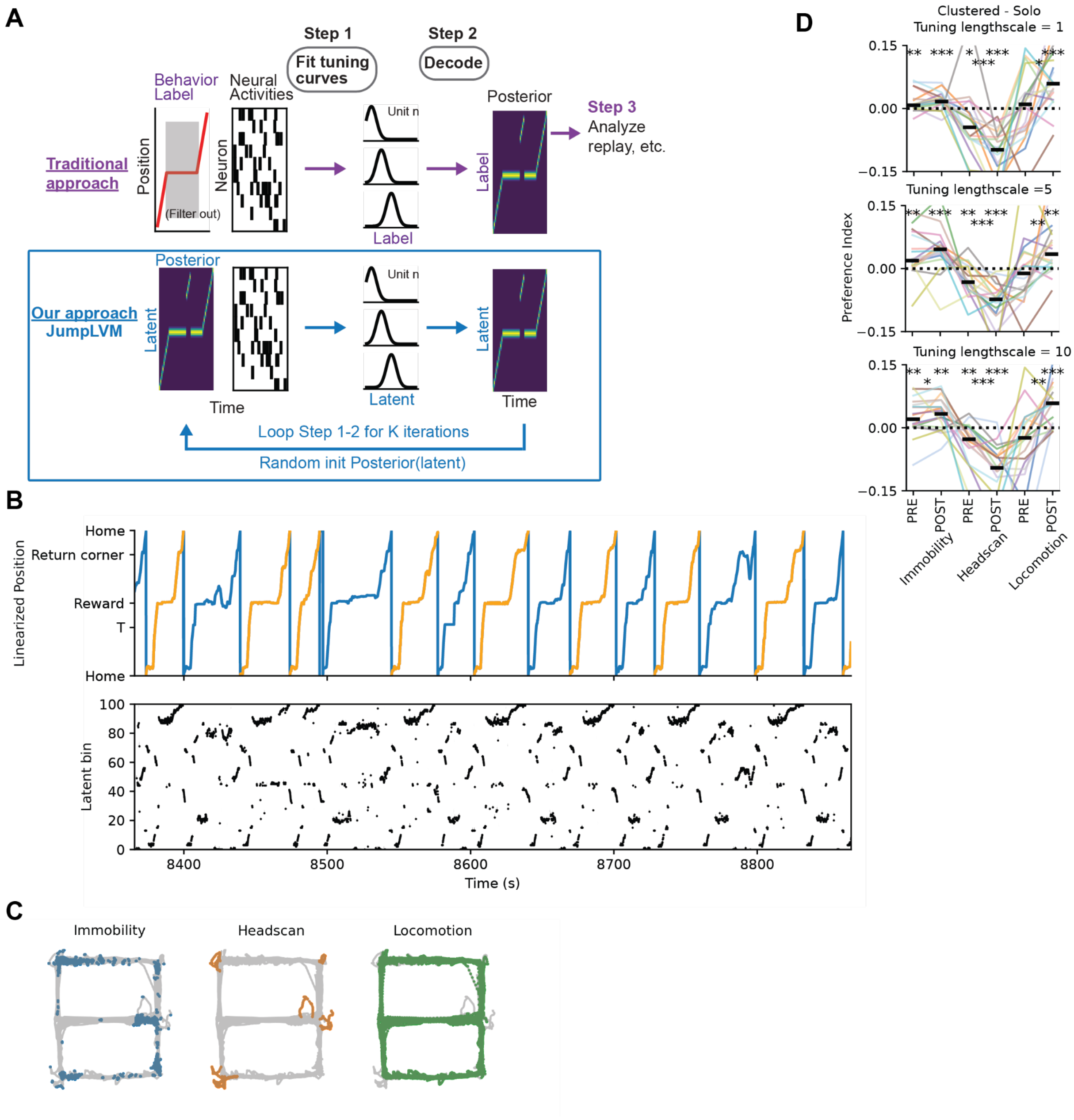


**Figure S13. Description of JumpLVM and behavior classification.** (**A**) Flowcharts of traditional and JumpLVM approaches. Traditional approach (purple arrows) starts with (filtered) behavior label (template) and neural activities, fits tuning curves, decodes and analyzes the decoded results. The JumpLVM approach (blue arrows) starts with random (but can be label-based) initialization (init) of the latent posterior and neural activities, and iteratively fits tuning curves and decodes and, crucially, uses the result of decoding as the input for the next iteration of tuning curve fitting. (**B**) Top: linearized position as the animal performs an alternation task in the figure-eight maze. Color indicates the turn performed in that trial. Bottom: posterior probability of the latent variable from the trained JumpLVM. Darker color means higher probability. (**C**) Each dot marks the position of the animal on the maze during each bin classified as Immobility (left), Headscan (middle) or Locomotion (right). (**D**) Clustered-Solo Preference Index per latent bin, separated by epoch (PRE-task sleep or POST-task sleep) and latent types. Each panel is the model fitted with a different tuning length scale. Each colored line is a session. Solid black short lines are the medians across sessions. The dotted black line indicates no preference for **c**SPW-Rs or **s**SPW-Rs. Headscan latents prefer **s**SPW-Rs, while locomotion latents gain preference for **c**SPW-Rs. Significance was determined via Wilcoxon rank-sum tests between PRE and POST, and via Wilcoxon signed-rank tests for each preference index against zero. The pattern is robust across different model hyperparameters.

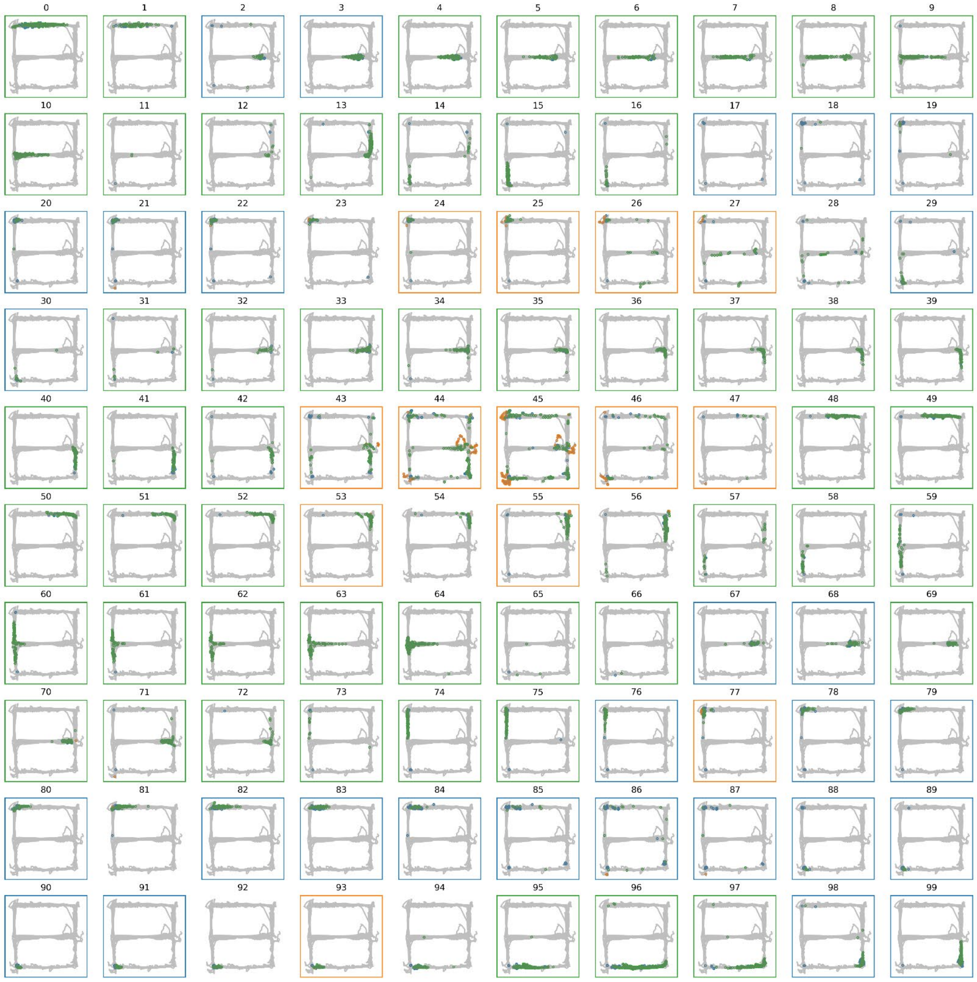


**Figure S14. Activation patterns of all latent bins in one session from JumpLVM.** Each panel is one latent bin: each dot marks the 2D position of the animal when that latent bin achieves MAP, colored by the behavior type of that time (blue: Immobility, orange: Headscan, green: Locomotion). The colored boxes indicate the classified type of that latent bin. Notice that the classification is based on probability and the dot here is only placed if the latent achieves maximum probability.

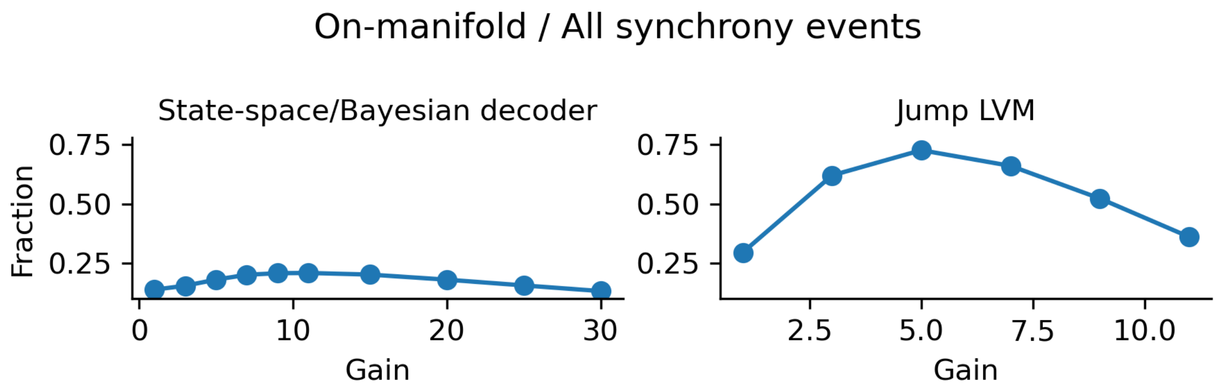


**Figure S15. Demonstration of the gain selection for using JumpLVM to decode synchrony events (example session).** Y axis: fraction of on-manifold synchrony events for state-space decoder (left) and JumpLVM (right). X axis: gain to be multiplied to the tuning curves before the decoding. For JumpLVM (right), the gain that maximized the fraction (on a random half of the events) was selected. For the state-space decoder the change in the fraction from changing the gain is small, so we used the default of 1. For all gain values JumpLVM finds more on-manifold events. (For the state-space decoder, we used a version of the marginal likelihood ignoring the temporal transition here to make it comparable to how we computed the marginal likelihood for JumpLVM, and thus it is the same as the marginal likelihood of a Naive Bayes decoder.)

**Supplementary Table 1**

Summary of recorded putative single units. We recorded single unit activity across CA3, CA1 and RSC. Neurons were classified as putative pyramidal cells (PYR) and putative interneurons (INT) based on their waveform and spiking statistics.

| Session Name | CA1 PYR | CA1 INT | CA3 PYR | CA3 INT | RSC PYR | RSC INT |
|---|---|---|---|---|---|---|
| TES_sResp_M01_20240313 | 85 | 27 | 31 | 18 | 56 | 23 |
| TES_sResp_M01_20240314 | 67 | 28 | 36 | 26 | 57 | 28 |
| TES_sResp_M02_20240312 | 63 | 18 | 7 | 17 | 3 | 2 |
| TES_sResp_M02_20240313 | 173 | 37 | 54 | 46 | 11 | 10 |
| TES_sResp_M02_20240314 | 156 | 31 | 60 | 55 | 13 | 18 |
| TES_sResp_M02_20240318 | 78 | 19 | 19 | 20 | 6 | 8 |
| TES_sResp_M03_20240621 | 91 | 39 | 61 | 13 | 17 | 26 |
| TES_sResp_M03_20240622 | 88 | 42 | 51 | 12 | 13 | 31 |
| TES_sResp_M03_20240623 | 133 | 51 | 70 | 27 | 19 | 19 |
| TES_sResp_M03_20240624 | 123 | 44 | 62 | 17 | 19 | 17 |
| TES_sResp_M05_20240729 | 72 | 128 | 100 | 16 | 21 | 35 |
| TES_sResp_M05_20240730 | 44 | 113 | 67 | 13 | 15 | 26 |
| TES_sResp_M05_20240731 | 57 | 141 | 55 | 17 | 15 | 32 |
| TES_sResp_M05_20241014 | 31 | 95 | 32 | 10 | 8 | 15 |

**Supplementary Table 2**

Statistical comparison of solo and cluster SPW-R properties during Wake and NREM states. R1, R2, and R3 refer to the first, second and third ripples within a cluster event. All values are FDR-corrected p-values; significant values ($p < 0.05$) are in bold.

| **Brain State** | **Wake** | | | **NREM** | | |
|---|---|---|---|---|---|---|
| **Ripple property** | **Duration (ms)** | **Amplitude (μV)** | **Peak frequency (Hz)** | **Duration (ms)** | **Amplitude (μV)** | **Peak frequency (Hz)** |
| **Solo vs R1** | 2.14e-01 | 2.24e-01 | 5.91e-01 | 3.97e-01 | **5.12e-06** | 3.03e-01 |
| **Solo vs R2** | 2.69e-01 | **4.68e-03** | 4.76e-01 | **1.66e-03** | **4.95e-22** | **5.54e-05** |
| **Solo vs R3** | **2.66e-02** | **4.68e-03** | 5.91e-01 | **1.79e-13** | **8.31e-15** | **9.72e-08** |
| **R1 vs R2** | 6.75e-01 | 7.79e-02 | 5.91e-01 | **1.10e-04** | **4.91e-12** | **1.13e-06** |
| **R2 vs R3** | 2.69e-01 | 7.79e-02 | 7.49e-01 | **5.32e-15** | **2.50e-05** | **1.63e-09** |
| **R1 vs R3** | 2.14e-01 | 9.27e-01 | 5.91e-01 | **6.47e-07** | **5.92e-04** | 1.13e-01 |

**Supplementary Table 3**

Summary statistics of spiking activity in CA1, CA3 and RSC during solo and cluster SPW-Rs.

| | **CA1 PYR** | **CA3 PYR** | **RSC PYR** | **CA1 INT** | **CA3 INT** | **RSC INT** |
|---|---|---|---|---|---|---|
| **Solo vs R1** | 0.313 | 0.053 | 0.348 | 0.878 | <0.001 | 0.172 |
| **Solo vs R2** | 0.055 | 0.239 | 0.025 | 0.321 | 0.041 | 0.172 |
| **Solo vs R3** | 0.055 | 0.037 | <0.001 | 0.017 | 0.644 | 0.051 |
| **R1 vs R2** | 0.367 | 0.472 | 0.159 | 0.329 | 0.06 | 0.968 |
| **R2 vs R3** | 0.856 | 0.314 | 0.001 | 0.026 | 0.017 | 0.419 |
| **R1 vs R3** | 0.389 | 0.683 | <0.001 | 0.017 | <0.001 | 0.419 |
